# The blood vasculature instructs lymphatics patterning in a SOX7 dependent manner

**DOI:** 10.1101/2021.07.02.450967

**Authors:** Ivy Kim-Ni Chiang, Winnie Luu, Key Jiang, Nils Kirschnick, Mehdi Moustaqil, Tara Davidson, Emmanuelle Lesieur, Renae Skoczylas, Valerie Kouskoff, Jan Kazenwadel, Luis Arriola-Martinez, Emma Sierecki, Yann Gambin, Kari Alitalo, Friedmann Kiefer, Natasha L. Harvey, Mathias Francois

## Abstract

Despite a growing catalogue of secreted factors critical for lymphatic network assembly, little is known about the mechanisms that modulate the expression level of these molecular cues in blood vascular endothelial cells (BECs). Here, we show that a BEC-specific transcription factor, SOX7, plays a crucial role in lymphatic vessel patterning by modulating the transcription of lymphangiocrine signals. While SOX7 is not expressed in lymphatic endothelial cells (LECs), loss of SOX7 function in mouse embryos causes a dysmorphic dermal lymphatic phenotype. We identify novel distant regulatory regions in mice and humans that contribute to directly repressing the transcription of a major lymphangiogenic growth factor (*Vegfc*) in a SOX7-dependent manner. Further, we show that SOX7 directly binds HEY1, a canonical repressor of the Notch pathway, suggesting that transcriptional repression may also be modulated by the recruitment of this protein partner at *Vegfc* genomic regulatory regions. Our work unveils a role for SOX7 in modulating downstream signalling events crucial for lymphatic patterning, at least in part via the transcriptional repression of VEGFC levels in the blood vascular endothelium.

## Introduction

The long-standing observation of parallel networks between arteries and lymphatic vessels has hinted at a role for the blood vessels in guiding lymphatic patterning during embryogenesis (Kampmeier, 1969). This influence of the blood vasculature on the trajectory of lymphatic vessels has been shown using both zebrafish and mouse model systems (Bussmann et al., 2010; Vaahtomeri et al., 2017). It has been established that blood endothelial cells (BECs) produce lymphangiogenic guidance cues such as chemokine receptor 4/chemokine ligand 12, adrenomedullin and semaphorin3G (Cha et al., 2012; Liu et al., 2016; Uchida et al., 2015). In addition to the blood vasculature, other tissues such as the mesenchyme and immune cells play a critical role in informing lymphatic vessel assembly (Cahill et al., 2021; Gordon et al., 2010; Karkkainen et al., 2004), mostly via the production of vascular endothelial growth factor C (VEGFC) (Rauniyar et al., 2018). During the early steps of lymphangiogenesis, VEGFC from the mesenchyme stimulates the exit of specified lymphatic endothelial cell (LEC) progenitors from the cardinal veins (CVs) and inter-somitic veins, and a subsequent dorso-lateral migration to form the lymphatic plexus (Karkkainen et al., 2004). Loss of VEGFC function results in the failure of LECs to form a primary vascular network, and a disorganised accumulation of LEC progenitors within the walls of the CVs.

Despite a plethora of information on the mode of action, maturation process and signalling mechanisms of VEGFC, there is a lack of understanding of how this growth factor is transcriptionally regulated, especially in the endothelial compartment. While the hematopoietically expressed homeobox (HHEX) genetic pathway has been implicated as an endothelial-specific regulator upstream of VEGFC expression, only the BTB and CNC homology (BACH) family of transcription factors have been shown to directly transactivate the VEGFC promoter and these experiments were performed in human ovarian carcinoma ES2 cells (Cohen et al., 2020; Gauvrit et al., 2018; Schuermann et al., 2015). Thus, there is still a paucity of information on the molecular mechanisms that drive *Vegfc* transcription in the endothelial cell context, despite the fact that it is well established that endothelial-derived VEGFC is critical for blood vessel angiogenesis and coronary artery formation (Cao et al., 1998; Chen et al., 2014).

The SOX group F (SOXF) transcription factors (SOX7, SOX17 and SOX18) are known to play a conserved role in vascular development (François et al., 2010; Yao et al., 2019). It has been reported that SOXF factors interact genetically with both VEGFR2 and VEGFD to create a positive feedback loop during blood vascular angiogenesis (Duong et al., 2013; Kim et al., 2016). In these studies, VEGF was shown to enhance SOX7 and SOX17 expression, and modulate the nuclear localisation of the SOXF proteins. SOX7 and SOX17, in turn, up-regulate the expression of VEGF receptor, VEGFR2. In addition, SOXF proteins function directly upstream of the Notch pathway by modulating the activities of the *Dll4* and *Notch1* enhancers (Chiang et al., 2017; Sacilotto et al., 2013). Taken together, these lines of evidence established a major regulatory axis between SOXF, VEGFs and the Notch pathway, most likely at play during both arterio-venous specification and angiogenesis. However, it remains elusive whether this signalling axis is also involved in other vascular remodelling processes.

Here, we show that SOX7 plays a pivotal role in controlling lymphatic vessel morphogenesis, at least in part via a repressive mechanism of *Vegfc* transcription in the blood endothelium. Endothelial-specific loss of SOX7 function during the early steps of lymphatic specification in mouse embryos results in both an ectopic distribution of PROX-positive LEC progenitors in the medio-ventral aspects of the CVs, and disruption of dermal lymphatics vessel morphogenesis, as characterised by an excessive emergence of endomucin (EMCN)-derived LEC progenitors. Consistent with a role for SOX7 in transcriptional repression of *Vegfc*, we demonstrated an up-regulation of BEC-derived *Vegfc* upon depletion of SOX7 in both *in vivo* and *in vitro* model systems. Genome-wide analysis of SOX7 binding locations *in vivo* revealed that this transcription factor directly binds to putative repressive distal regulatory regions upstream of VEGFC transcription start sites, in both human and mouse. Finally, we showed that SOX7 physically interacts with the Notch effector and transcriptional repressor, HEY1, suggesting indirect transcriptional repression of *Vegfc* mediated by Notch signalling. This study identifies SOX7 as the first direct endothelial-specific regulator of *Vegfc* transcription, and uncovers novel mechanisms that likely underlie the SOX7-mediated fine-tuning of VEGFC expression in BECs, a process essential for correct lymphatic growth and patterning.

## Results

### The SOX7 BEC-specific transcription factor is necessary for dermal lymphatic patterning

While it is known that blood vessels instruct the patterning of lymphatic vessels in a paracrine manner, little is known about how the expression levels of guiding molecules from blood vessels are transcriptionally regulated. To address this, we took advantage of the loss-of-function of the mouse SOX7 transcription factor, which expresses specifically in blood endothelial cells (BECs), but not in lymphatic endothelial cells (LECs) (Kim et al., 2016). The endogenous expression profile of SOX7 was determined using a *Sox7*-V5 tagged transgenic reporter mouse line (Sup. Fig. 1). Briefly, the V5 epitope was knocked in within the C-terminal domain of *Sox7* to specifically label endogenous SOX7 expression. *Sox7*-V5 homozygous animals are fertile and viable. The *Sox7*-V5 tagged protein was detected specifically in endothelial cell (EC) nuclei of both the cardinal veins (CVs) and the dorsal aortae (DA), but none was observed in migrating LECs (Sup. Fig. 2A, B), consistent with a previous report (Kim et al., 2016). While SOX7 was detected in some ECs of the CVs (Sup. Fig. 2A, B), none was observed in mature veins and venules in embryonic skin (Sup. Fig. 2C). High levels of *Sox7*-V5 were observed in arterioles near the midline but only low levels of SOX7 were detected in mature SOX17-positive arteries (Sup. Fig. 2C). These observations were further supported by immunofluorescence analysis for *β*-galactosidase (*β*-gal) in E14.5 embryonic skin of a *Sox7 lacZ* knock-in mouse line (Sup. Fig. 2D).

**Figure 1.**
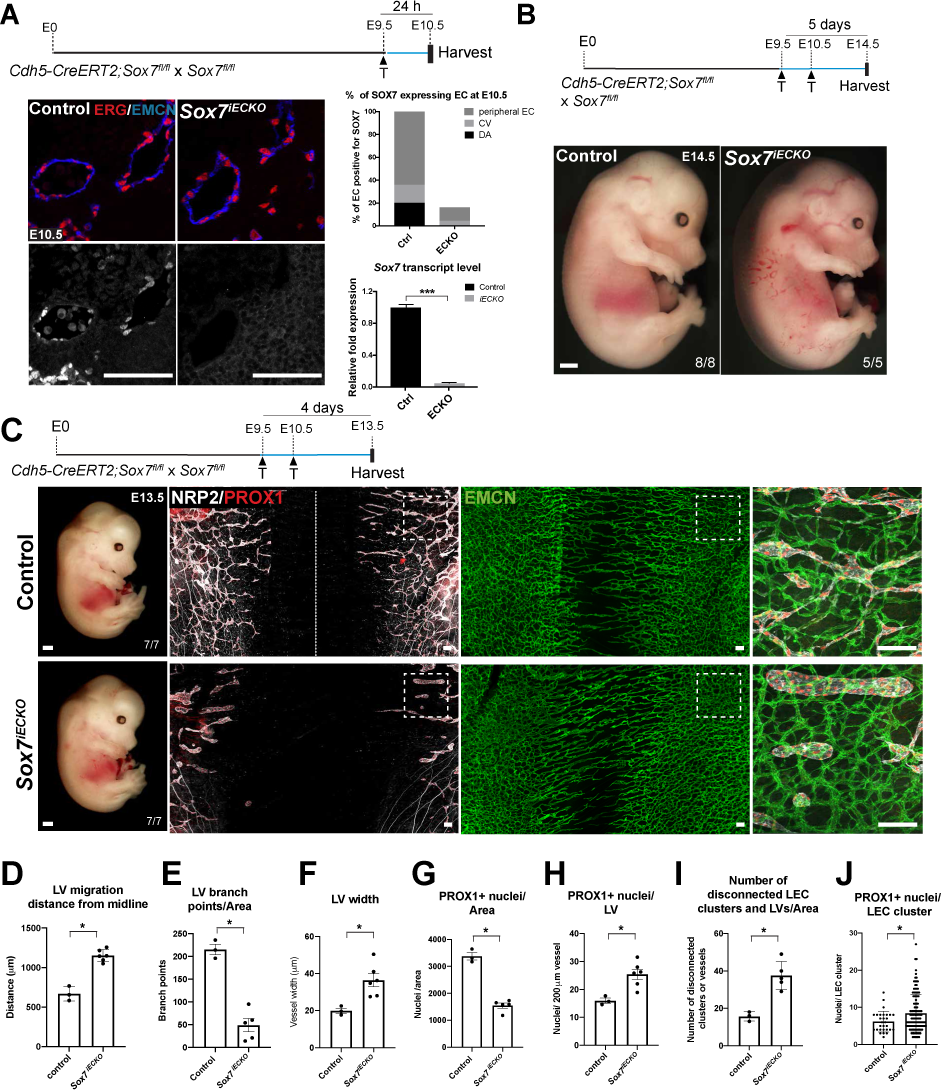
Conditional loss of *Sox7* function in blood vascular endothelial cell causes dermal lymphatic patterning defects. (A) Representative images of E10.5 *Sox7^iECKO^* mutants and sibling controls, injected with tamoxifen at E9.5. The endothelium is delineated by endomucin (EMCN) (blue) and endothelial cells by ETS-related gene (ERG) (red). (Top right panel) The percentage of SOX7 positive endothelial cells over the total number of endothelial cells quantified from 10 sequential transverse sections in 2 control and 3 *Sox7^iECKO^* mutants. (Bottom right panel) Graph indicating the levels of *Sox7* transcript in sibling controls and *Sox7^iECKO^* mutants. Mean ± SEM; scored sibling control, n=9; *Sox7^iECKO^* mutants, n=7; Mann-Whitney *U-*test. *P*<0.0005 (***). **(B)** Bright-field images of *Sox7^iECKO^* mutants and sibling controls at E14.5, after pulsing with tamoxifen at E9.5 and E10.5. Scored sibling controls, n=8; *Sox7^iECKO^* mutants, n=5. **(C)** Brightfield images and whole-mount immunostaining of *Sox7^iECKO^* mutant and sibling control skin at E13.5, after Cre induction at E9.5 and E10.5. Dermal lymphatic structures are marked by Neuropilin 2 (NRP2) (membranous white), lymphatic endothelial cells by Prospero-related homeobox 1 (PROX1) (red), and blood vessels by EMCN (green). Dash line represents the midline of the embryo. Scored sibling controls, n=7; *Sox7^iECKO^* mutants, n=7 **(D-H)** Quantification of **(D)** lymphatic sprout migration distance from the midline **(E)** lymphatic branch points/area **(F)** lymphatic vessel width (μm) **(G)** PROX1+ nuclei/area and **(H)** PROX1+ nuclei/lymphatic vessels across the whole skin. Total area = 4200 x 1500 μm on both sides of the midline, in sibling controls (n=3) and *Sox7^iECKO^* mutants (n=5-6). **(F,H)** Average of width and PROX1+ nuclei was obtained from 7 random representative leading lymphatic vessels, of fixed length at 200 μm, from both sides of the midline in each skin. **(I,J)** Quantification of **(I)** disconnected lymphatic endothelial cell (LEC) clusters (<100 μm) and vessel branches (>100μm)/area and **(J)** PROX1+ nuclei in each LEC cluster. Total area quantified = 4200 x 2200 μm on both side of midline in sibling controls (n=3) and *Sox7^iECKO^* mutants (n=5). PROX1+ nuclei were quantified from n=29 and n=122 LEC clusters, respectively. **(D-J)** Skins were from the cervico-thoracic regions of E13.5 embryos, defloxed at E9.5, E10.5. Mean ± SEM, Mann-Whitney *U*–test. *P*<0.05 (*). LV, lymphatic vessel. Scale bars = 100 μm (immunofluorescence images A,C), 1 mm (bright-field images B,C).

**Figure 2.**
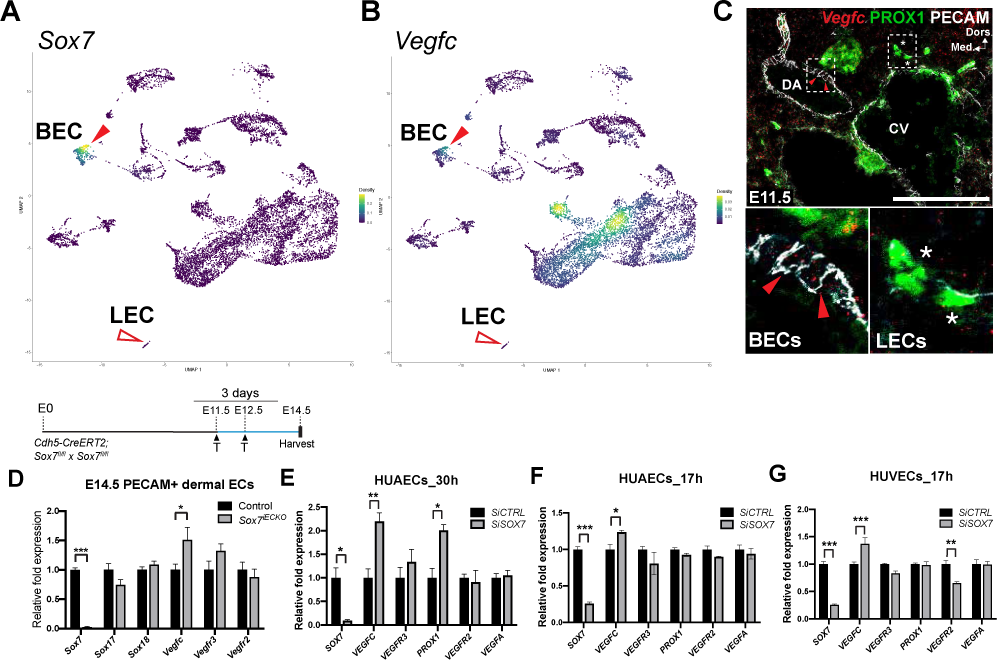
Loss of *Sox7* function upregulates *Vegfc* transcript levels in blood endothelial cells. (A-B) Nebulosa plots from single-nuclei RNA-Seq on E14.5 embryonic skins, showing the expression of *Sox7* and *Vegfc.* Both *Sox7* and *Vegfc* are found expressed in BECs (red arrowheads), but not LECs (empty red arrowheads). **(C)** *Vegfc* (red) fluorescent RNA *in situ* hybridisation on mouse cross-sections at E11.5. The endothelium is delineated by PECAM (white) and LECs by PROX1 (green nuclei). BEC-specific endogenous levels of *Vegfc* (red arrowheads) are higher than LECs (asterisks). **(D)** qPCR on FACs-sorted PECAM+CD45- endothelial cells of *Sox7^iECKO^* mutants and sibling controls at E14.5, injected with tamoxifen at E11.5 and E12.5. Expression is normalised to endothelial marker *Pecam* and *Cdh5*. Scored sibling controls, n=9; *Sox7^iECKO^* mutants, n=8. **(E-F)** qPCR on human arterial endothelial cells (HUAECs) transfected with *SiSOX7* or *SiCTRL* for **(E)** 30 h and **(F)** 17 h. **(G)** qPCR on human venous endothelial cells (HUVECS) transfected with *SiSOX7* or *SiCTRL* for 17 h. **(E-G)** Expression is relative to *HPRT* and *GAPDH.* Data from one siRNA experiment performed in triplicates. **(D–G)** Mean ± SEM; *t*-test. *P*<0.05 (*); *P*<0.005 (**); *P*<0.0005 (***). BEC, blood endothelial cell; LEC, lymphatic endothelial cell; DA, dorsal aorta; CV, cardinal vein; Dors., dorsal; Med., medial. Scale bars = 100 μm.

To explore the role of SOX7 in lymphatic vascular remodelling, we took advantage of a SOX7 loss-of-function mouse model. It has been shown that *Sox7^-/-^* homozygous mutants die *in utero* by E10.5 (Kim et al., 2016; Lilly et al., 2017), so we used an endothelial-specific Cre inducible mouse line, the *Cdh5-CreERT2:Sox7 fl/fl* mouse line. Tamoxifen-induced excision of SOX7 exon 2 was performed at E9.5 and E10.5, the stage of early LEC specification in the CVs (Srinivasan et al., 2007; Wigle and Oliver, 1999). Cre efficiency was initially validated by a single pulse of tamoxifen at E9.5, followed by analysis at E10.5 (Fig. 1A). Further analysis followed tamoxifen doses at E9.5 and E10.5. At E14.5, the Cre-inducible endothelial-specific *Sox7* knockout (*Sox7^iECKO^*) mutant embryos appeared similar in size to their sibling controls, but displayed gross subcutaneous edema, with some showing blood-filled lymphatic vessels (Fig. 1B, Sup. Fig. 3, 80% of mutant with edema out of n=5). While gross edema is less obvious a day earlier at E13.5 (Fig. 1C), detailed analysis of dermal lymphatics revealed that lymphatic vessels in the *Sox7^iECKO^* mutants were severely dilated, lacked inter-vessel branches and showed migration defects (Fig. 1C-F). Although there is a decrease in total PROX1+ cells in the skin from the flank (Fig. 1G), quantitative analysis of PROX1+ nuclei normalised to LEC branches across the entire skin sample (flank + midline) showed a significant increase in the mutant (Fig. 1H, J), suggesting LEC hyper-proliferation coupled to a failure of vessel remodelling. Relative to sibling controls, most LECs in the *Sox7^iECKO^* mutant embryos assemble in islands or clusters that appeared disconnected from the main network (Fig. 1C, I). Consistent with the gross edema observed at E14.5, the dermal lymphatic phenotype persists, and appeared further aggravated at this stage (Sup. Fig. 4A). Unlike the lymphatic vessels, blood vessel morphogenesis is less severely affected in E13.5 *Sox7^iECKO^* mutant embryos (Fig. 1C). Apart from increased blood vessel density at the angiogenic front, which is accompanied by an increase in the proportion of proliferative ERG+ endothelial cells (Sup. Fig. 4B); overall the blood vessel network in *Sox7^iECKO^* mutant embryos appeared intact and comparable to sibling controls. To rule out the possibility that the observed lymphatic patterning defect is due to *CreERT2* toxicity (Brash et al., 2020), we created *Cdh5-CreERT2: Sox7 fl/fl*; mT/mG and *Cdh5-CreERT2*; mT/mG animals. The mT/mG double fluorescence Cre-reporter mice express the tdTomato red fluorescent protein prior to Cre-mediated excision, and green fluorescent protein after excision, therefore allowing *Cdh5-CreERT2* activity to be measured and traced (Muzumdar et al., 2007). To effectively quantify *Cdh5-CreERT2* activity, mice were pulsed with tamoxifen at E12.5, and embryos harvested at E14.5. Despite showing comparable *CreERT2* activity to sibling controls (Sup. Fig. 4D-F), *Sox7^iECKO^; mT/mG* embryos consistently showed dilated lymphatic vessels with reduced branching points (Sup. Fig. 4C). This suggests that the mis-patterning of dermal lymphatics observed in *Cdh5-CreERT2: Sox7 fl/fl* embryos is dependent on SOX7 loss-of-function. Together, these results show that tightly regulated BEC-specific SOX7 expression is essential for correct patterning of the lymphatic network.

**Figure 3.**
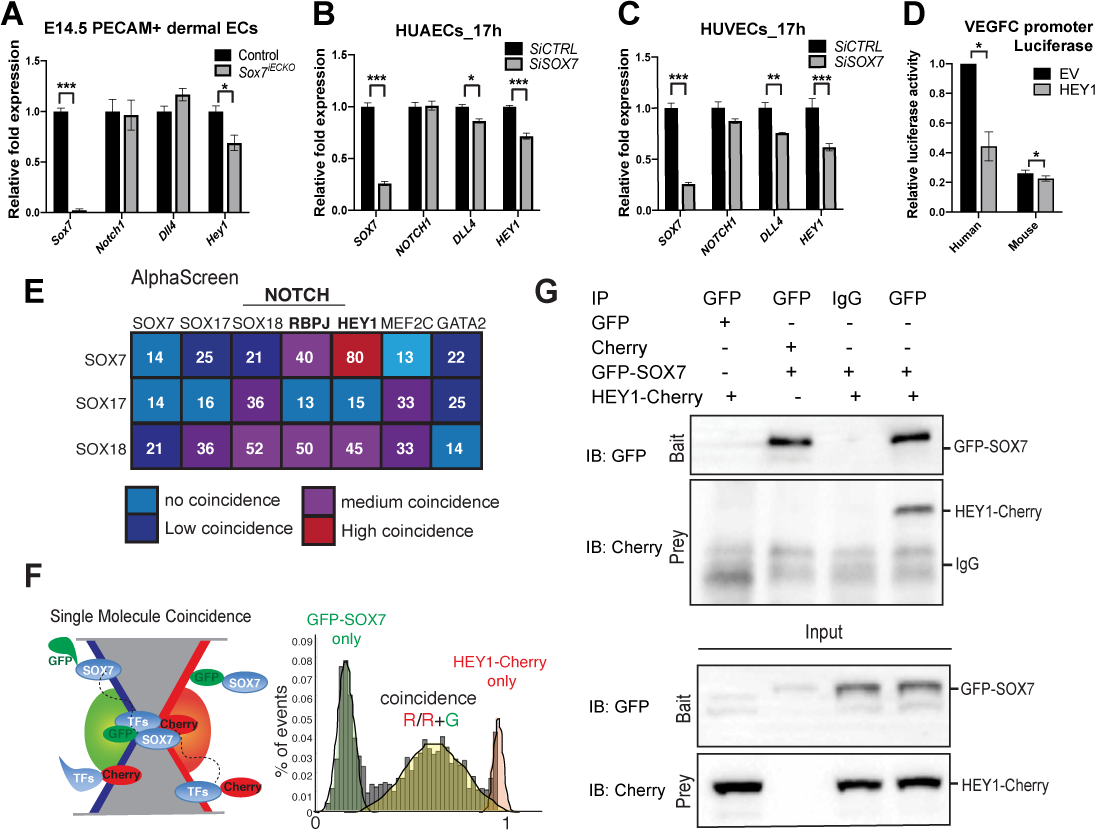
SOX7 indirectly suppresses VEGFC expression through the Notch pathway. (A-D) SOX7 transcriptionally activates Notch effector, HEY1, to repress *Vegfc*. **(A)** qPCR on FAC-sorted PECAM+CD45- endothelial cells of *Sox7^iECKO^* mutants and sibling controls at E14.5, injected with tamoxifen at E11.5 and E12.5. Expression is normalised to the endothelial marker *Pecam* and *Cdh5*. Scored sibling controls, n=9; *Sox7^iECKO^* mutants, n=8. **(B-C)** qPCR on **(B)** human arterial endothelial cells (HUAECs) and **(C)** human venous endothelial cells (HUVECs) transfected with *SiSOX7* or *SiCTRL* for 17 h. In addition to *HEY1, DLL4* levels were also downregulated in the human cell line experiments. Expression is relative to *HPRT* and *GAPDH.* Data from one siRNA experiment performed in triplicates. Mean ± SEM; *t*-test. *P*<0.05 (*); *P*<0.005 (**); *P*<0.0005 (***). **(D)** HEY1 represses human VEGFC promoter activity. HeLa cells were co-transfected with human or mouse VEGFC-luc and either EV (empty vector) or HEY1 expression constructs as indicated. VEGFC luciferase activity was measured and normalised to Renillla luciferase activity, which was then made relative to the promoter-less vector, pGL3-basic, which was set to 1. Biological replicates, n=3 independent repeats of the same experiment. Mean ± SEM; *t*-test. *P* <0.05 (*). **(E-G)** SOX7 physically interacts with transcription repressor, HEY1. **(E)** Amplified Luminescent Proximity Homogenous Assay (ALPHAScreen) shows the heatmap of SOX7 pairwise protein-protein interaction tested, where red indicates strong interaction and light blue indicates an absence of interaction. **(F)** Single molecule spectroscopy reveals that SOX7 is able to directly interact with HEY1. SOX7 and the target transcription factors were tagged with GFP or Cherry. GFP-SOX7 and HEY1-Cherry show co-incidence at 0.66, suggesting a 1:2 interaction. Peaks centered on 0 (green) or 1 (red) correspond to the GFP or Cherry-tagged proteins only. **(G)** Co-immnunoprecipitation analysis of SOX7 and HEY1. HEK293 cells were transfected with indicated plasmids and harvested 24 h after transfection, immunoprecipitated by anti-GFP or Ig control before immunoblot analysis to determine the presence of bait/prey (top) and the input (bottom). N=2.

**Figure 4.**
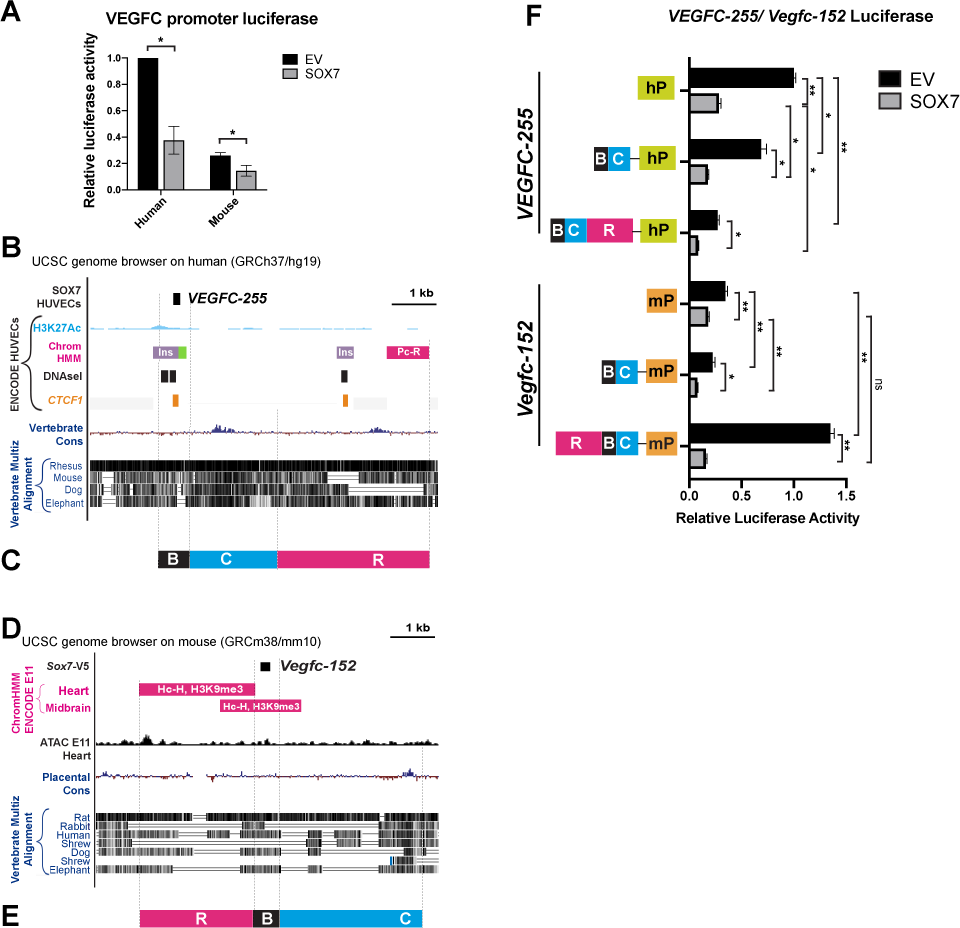
SOX7 directly binds to silencing regions at the *Vegfc* locus and represses *Vegfc* transcription. (A) SOX7 represses human VEGFC promoter activity. HeLa cells were co-transfected with human or mouse VEGFC-luc, EV (empty vector), or SOX7 expression constructs as indicated. VEGFC luciferase activity was measured and normalised to Renillla luciferase activity, which was then made relative to the promoter-less vector, pGL3-basic, which was set to 1. Biological replicates, n=3. Mean ± SEM; *t*-test. *P*<0.05 (*). **(B)** Schematic representation of the human *VEGFC* locus 255 kb upstream from the transcription start site (TSS) (denoted as *VEGFC-255* region) from the UCSC Genome browser. The H3K27Ac is denoted in light blue, DNAseI hypersensitive hotspots are indicated by black/grey boxes, where the darkness is proportional to the maximum signal strength observed in any cell line. The chromatin state in HUVECs is shown in purple (indicates insulator), green (poised enhancer) and pink (a Polycomb-repressed region). The HUVECs CCCTC-binding factor (CTCF) binding location is shown in orange. Multiple species conservation is shown in blue peaks and alignments with black stripes. **(C)** The human luciferase *VEGFC-255 BCR* transgene. B (black), binding region; C (blue), conserved region and R (pink), repressive region. **(D)** Schematic representation of the mouse *Vegfc* locus 152kb upstream from the TSS (denoted as *Vegfc-*152). The chromatin state in mouse at E11 from ENCODE is denoted in pink (indicates a heterochromatin region enriched for H3K9me3 repressive marker). The ATAC-Seq (in black) indicates accessible DNA regions in E11 mouse hearts. Multiple species conservation is shown in blue peaks and alignments with black stripes. **(E)** The mouse luciferase *Vegfc-152 RBC* transgene. R (pink), repressive region; B (black), binding region and C (blue), conserved region. **(F)** SOX7 represses VEGFC promoter activity through *VEGFC-255* and *Vegfc-152*. HeLa cells were co-transfected with human or mouse VEGFC-luc, *VEGFC-255* BC-luc, *VEGFC-255* BCR-luc, *Vegfc-152* BC-luc, *Vegfc-152* RBC-luc, EV (empty vector), or SOX7 expression constructs as indicated. VEGFC luciferase activity was measured and normalised to Renillla luciferase activity, which was then made relative to the promoter-less vector, pGL3-basic, which was set to 1. Biological replicates, n=3. Mean ± SEM; *t*-test. *P* <0.05 (*); *P*<0.005 (**); ns = not significant.

### SOX7 is a negative regulator of VEGFC transcription in blood vascular endothelial cells

The expression profile of SOX7 in angiogenic blood vessels (Sup. Fig. 2C) (del Toro et al., 2010), combined with the observation of lymphatic patterning defects in SOX7 loss-of-function mouse embryos, prompted us to investigate the levels of lymphangiogenic molecules in BECs. To profile lymphangiocrine signals in BEC during development, we performed single-nuclei RNA-Seq on whole skin from wild-type mouse embryos at E14.5. This approach reveals that only BECs and a sub-population of neurons and smooth muscle cells actively transcribe *Vegfc* during dermal embryogenesis. Consistent with the immunofluorescence analysis in Sup Fig 2B, we observed *Sox7* expression restricted to endomucin-positive BECs with no expression in *Prox1*-positive LECs or other non-endothelial cell types (Fig. 2A, Sup. Fig. 5A-D). Interestingly, no *Vegfc* transcripts were detected in *Prox1*-positive LECs (Fig. 2B and Sup. Fig. 5B); suggesting therefore VEGFC autocrine signalling is not at play in dermal LECs. Co-localisation between *Sox7* and *Vegfc* expression in a subset of endomucin-positive BECs raises the possibility that SOX7 may transcriptionally regulate *Vegfc* in this vascular plexus (Fig. 2A, B, red arrowheads). We then sorted BECs and LECs from E14.5, E16.5 and E18.5 mouse embryonic skins (Kazenwadel et al., 2012) and performed microarray analysis on both cell populations. This approach confirmed that during development, BECs express *Vegfc* mRNA at 1.5-2.9-fold higher levels than LECs (Sup. Fig. 5E). This result was further validated in mouse by *Vegfc* single molecule fluorescent *in situ* hybridisation (ISH) using RNAscope analysis in wild-type embryos at around E11.5 (Fig. 2C). Thick sections of wild-type embryos were labelled for *Vegfc* (red), PECAM (white) and PROX1 (green), demonstrating that the BEC-specific endogenous levels of VEGFC (Fig. 2C, red arrowheads) are higher than LECs (Fig. 2C, white asterisks). VEGFC expression in BECs has been observed previously in developing mouse and zebrafish DA, by immunostaining and *in situ* hybridisation assay, respectively (Hogan et al., 2009; Karkkainen et al., 2004).

**Figure 5.**
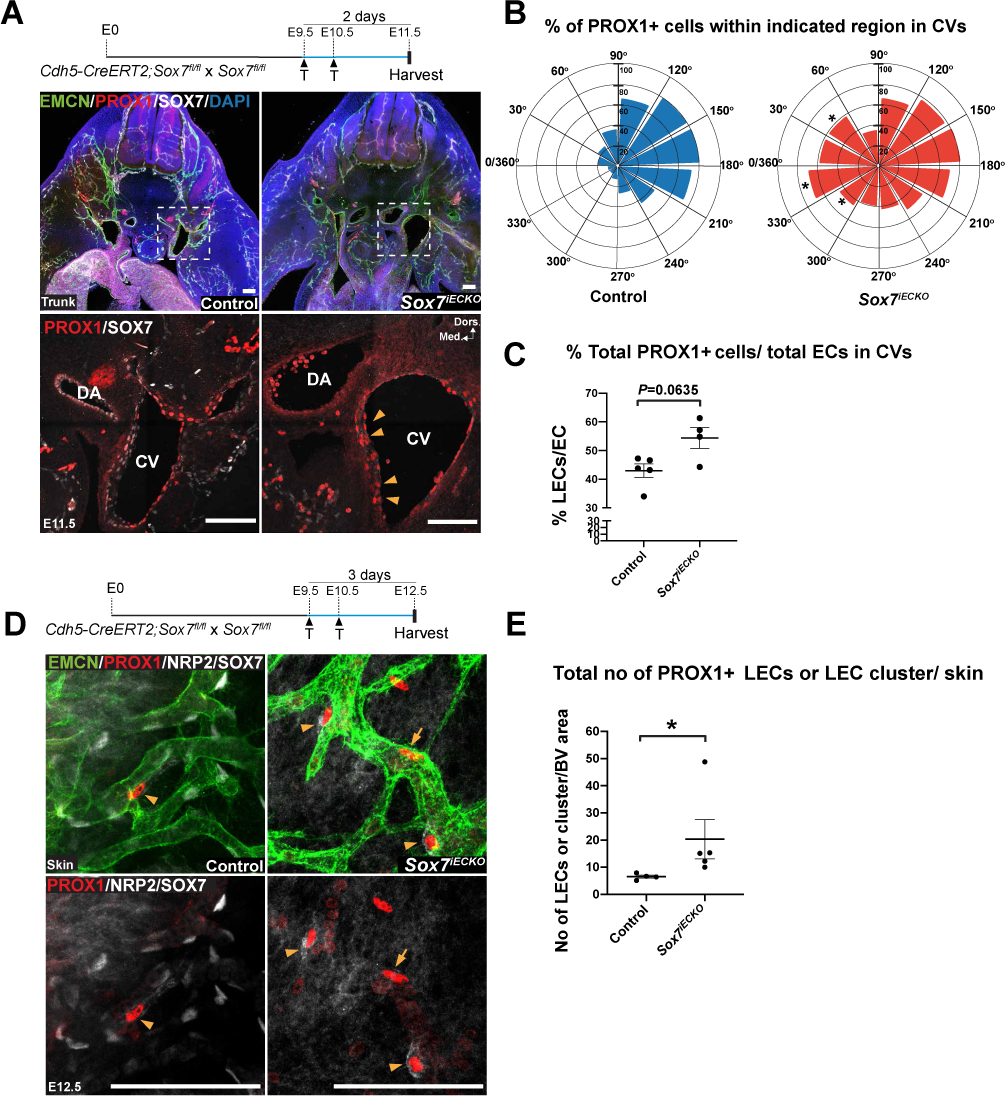
Loss of SOX7 transcription factor function causes ectopic LEC localisation in the cardinal veins and an excess of local LEC progenitors in embryonic skin. (A) At 2 days post Cre induction, E11.5 *Sox7^iECKO^* mutants showed ectopic expression of PROX1+ (red) in medial ventral regions of the cardinal veins (CVs), close to dorsal aorta (DA) (yellow arrowheads). **(B)** Rose diagram indicates the % of PROX1+ cells over total number of EMCN+ endothelial cells, within each indicated region. The medial region closest to the DA is indicated by 0/360°; most lateral region, 180°; dorsal, 90° and ventral, 270°. Scored sibling controls, n=5; *Sox7^iECKO^* mutants, n=4. Mean ± SEM; Mann-Whitney *U*-test. *P*<0.05 (*). **(C)** Graph indicates the % of total PROX1+ cells over the total number of EMCN+ endothelial cells within each embryo analysed. A total of 2907 endothelial cells quantified, from n=5 sibling controls and 1780 endothelial cells from n=4 *Sox7^iECKO^* mutants. Mean ± SEM; Mann-Whitney *U*-test. *P*=0.0635. **(D)** Whole-mount immunostaining of control and *Sox7^iECKO^* embryonic skin at E12.5 after *Cre* induction at E9.5, E10.5. Dermal lymphatic structures are stained with NRP2 (membranous white), lymphatic endothelial cells, PROX1 (red), and blood vessels, both EMCN (green) and SOX7 (white nuclei). Arrowheads indicate EMCN^low/-^PROX1^+^NRP2^+/low^ individual LECs on blood vessels (type I), and arrow shows EMCN^+^PROX1^+^ NRP2^low/-^ LEC within the vessel wall (type II). **(E)** Graph shows the total events of PROX1+ single LECs or LEC clusters cells associated with EMCN+ blood vessel plexus within each embryo. PROX1+ cells were quantified from n=4 controls and n=5 *Sox7^iECKO^* mutants. Mean ± SEM; Mann-Whitney *U*-test. P<0.05 (*). Dors., dorsal; Med., medial. Scale bars = 100 μm.

To assess whether the VEGF/VEGFR pathway is affected at a transcriptional level by loss of SOX7 function, we compared the expression of key genes by qPCR profiling of sorted CD31+CD45- endothelial cells from skin of E14.5 control and *Sox7^iECKO^* embryos, following tamoxifen induction at E11.5 and E12.5 (Fig. 2D, Sup. Fig. 4G). This analysis showed a significant up-regulation of *Vegfc* mRNA levels in *Sox7^iECKO^* mutant cells. To test whether SOX7 cell autonomously regulates *VEGFC* transcript levels in BECs, we next reduced levels of *SOX7* using a siRNA-based approach in human EC lines (HUAECs and HUVECs). At 30 h post-siRNA transfection, we observed a 2 fold-increase in the level of *VEGFC* transcripts in HUAECs (Fig. 2E). To reduce the number of potential secondary hits mediating the increase of *VEGFC* levels downstream of SOX7, gene expression analysis was performed at 17 h post-*SOX7* gene depletion. Both HUAECs and HUVECs showed a significant increase in *VEGFC* transcript levels in response to depleted SOX7 function (Fig. 2F, G). A significant reduction of *VEGFR2* transcripts was only observed in *SOX7* siRNA depleted HUVECs but not HUAECs, suggesting that transcriptional regulation of *VEGFR2* by SOX7 is context dependent. The consistent upregulation of *VEGFC* in response to *SOX7* gene depletion both *in vivo* and *in vitro* implies that *VEGFC* is an early response gene that is negatively regulated by SOX7 activity.

### SOX7 represses VEGFC indirectly through up-regulation of the Notch effector HEY1

Having established that BEC-specific expression of *VEGFC* is regulated by SOX7 activity, we next investigated how SOX7 might exert its repressive effects on *VEGFC* gene expression. The Notch signalling pathway has been previously shown to negatively-regulate specification of LECs in the ventro-medial aspects of the CVs, with ectopic Notch signalling reducing levels of the LEC specification regulator, PROX1 (Murtomaki et al., 2013). This finding, together with the known role of SOX7 in regulation of *Notch1/Dll4* transcription (Chiang et al., 2017; Sacilotto et al., 2013) lead us to postulate that SOX7 may negatively-regulate *Vegfc* through modulation of the Notch pathway. To evaluate the level of Notch activity in the context of SOX7 loss of function, we assessed the transcript levels of key Notch pathway members in sorted CD31+CD45- endothelial cells from E14.5 mouse skin samples, We observed a significant downregulation of the Notch effector, *Hey1*, but not *Dll4* and *Notch1,* in cells from *Sox7^iECKO^* embryos versus sibling controls (Fig. 3A). In siRNA experiments in human cells, at 17 h post-siSOX7 transfection levels of both *DLL4* and *HEY1* were consistently reduced in HUAECs and HUVECs (Fig. 3B, C).

HEY1 is a known transcriptional repressor that exerts its repressive activity through its basic helix-loop-helix (bHLH) domain (Fischer et al., 2005; Nakagawa et al., 2000). HEY1 has been shown to suppress the VEGFC receptor, VEGFR3 (Murtomaki et al., 2013). To test if HEY1 could also directly repress VEGFC, we overexpressed HEY1 in HeLa cells transfected with a synthetic construct containing a luciferase reporter gene fused to either a mouse or a human VEGFC promoter fragment (Chilov et al., 1997; Huang et al., 2014). In both cases, we found that HEY1 overexpression significantly reduced the VEGFC promoter activity (Fig. 3D), showing that HEY1 has the capacity to repress *Vegfc* transcription. Together, these observations support that SOX7 indirectly represses *Vegfc* through transactivation of the Notch effector and transcriptional repressor, HEY1.

### SOX7 interacts with transcriptional repressor, HEY1

In addition to directly regulating gene transcription, there is a growing body of evidence suggesting that SOXF factors modulate gene expression through protein-protein interactions (PPIs) with other transcription factors (Hosking et al., 2001; Lilly et al., 2016; Overman et al., 2017; Sacilotto et al., 2013). Given that Sox7 and Hey have been previously shown to genetically interact in the zebrafish vasculature (Hermkens et al., 2015), we investigated whether SOX7 physically interacts with HEY1 directly through PPIs.

To screen for PPIs in a low-throughput manner, we used a cell-free protein production system (Leischmania Tarentolae) combined with an Amplified Luminescent Proximity Homogenous (ALPHA) Screen assay. This approach enabled us to assess multiple pair-wise interactions between SOX7 and other SOXF members (e.g. SOX17 and SOX18), and key effectors of the Notch pathway such as RBPJ and HEY1, as well as other transcription factors such as GATA2. We used well-characterised PPIs such as SOX18/SOX18 homodimer and SOX18/RBPJ, or SOX18/MEF2C heterodimer (Moustaqil et al., 2018; Overman et al., 2017), as positive controls to calibrate the strength of the ALPHAScreen signal. An ALPHAScreen signal of medium and high coincidence was detected for SOX7/RBPJ and SOX7/HEY1 respectively, suggesting robust direct PPIs (Fig. 3E). To further validate interactions with the highest coincidence score, we took advantage of a single molecule two-colour coincidence assay (Fig. 3F) previously use to determine stoichiometric ratios for SOX18 protein partners (Moustaqil et al., 2018; Sierecki et al., 2014). In the case of the SOX7 and HEY1 interaction, the coincidence ratio in the same confocal volume of SOX7-GFP and HEY1-mCHERRY individual molecules was 0.5 (Fig. 3F, yellow trace). This ratio defines a stoichiometric relationship of 1:1. Further validation of SOX7/HEY1 protein recruitment was performed *in vitro* using co-immunoprecipitation experiments in HEK293 cells transiently transfected with epitope-tagged constructs (Fig. 3G). Together, we show that SOX7 is capable of physically engaging with HEY1, and raise the possibility that SOX7 not only indirectly represses *VEGFC* transcription through modulation of HEY1 levels, but can also coordinate transcriptional repression through a direct protein recruitment process at *VEGFC* regulatory regions on the genome.

### SOX7 directly represses VEGFC by acting on its promoter and distal silencing elements

To further investigate the possibility of an alternative mode of SOX7-dependent repression of *VEGFC* gene expression, we next explored the possibility of direct repression of *VEGFC* transcription by SOX7. Using the mouse and human VEGFC promoter luciferase reporter system described above, we assessed the baseline activity of both constructs either in the absence or presence of SOX7. Surprisingly, SOX7 dramatically suppressed both mouse and human VEGFC promoters (Fig. 4A), supporting the notion that SOX7 has the potential to act as a direct negative regulator of *VEGFC* transcription.

This observation suggested that SOX7 might interact with other *VEGFC* regulatory regions more broadly on a genome-wide scale. To probe for SOX7 genome binding locations, we took advantage of our SOX7-mCherrry ChIP-Seq dataset from human venous endothelial cells (HUVECs) (https://www.ebi.ac.uk/arrayexpress/experiments/E-MTAB-4480/) (Overman et al., 2017). Analysis of this dataset using the top 2k most intense ChIP-seq peaks revealed that the VEGF ligand-receptor interaction is one of the top 4 most enriched pathways (Sup. Fig. 6A), thereby confirming that SOX7 preferentially binds to regions associated to the VEGF/VEGFR pathways. We identified a putative *VEGFC* regulatory region situated 255 kb upstream from the VEGFC transcription start site (TSS), named *VEGFC*-255 (Fig. 4B). Although the *VEGFC-255* region does not overlap with either active (H3K27Ac) or repressive (H3K9me3, H3K27Me3) histone marks, it coincides with an open chromatin region (as revealed by DNAseI) where the transcription repressor and insulator CTCF binds (Fig. 4B) (Cuddapah et al., 2009; Filippova et al., 1996). Further, chromatin state segmentation analysis (Chrom HMM) from ENCODE confirmed that the binding region *VEGFC-255* has been associated with insulators (purple box) and poised enhancers (green box) (Fig. 4B). Interestingly, poised enhancers are usually bound by Polycomb Repressive Complex (Pc-R) (Cruz-Molina et al., 2017), and we observed a large Polycomb-repressed region situated around ∼5 kb upstream from *VEGFC*-255 (Pink box, Fig. 4B), which might suggest that this region has a repressive function.

**Figure 6.**
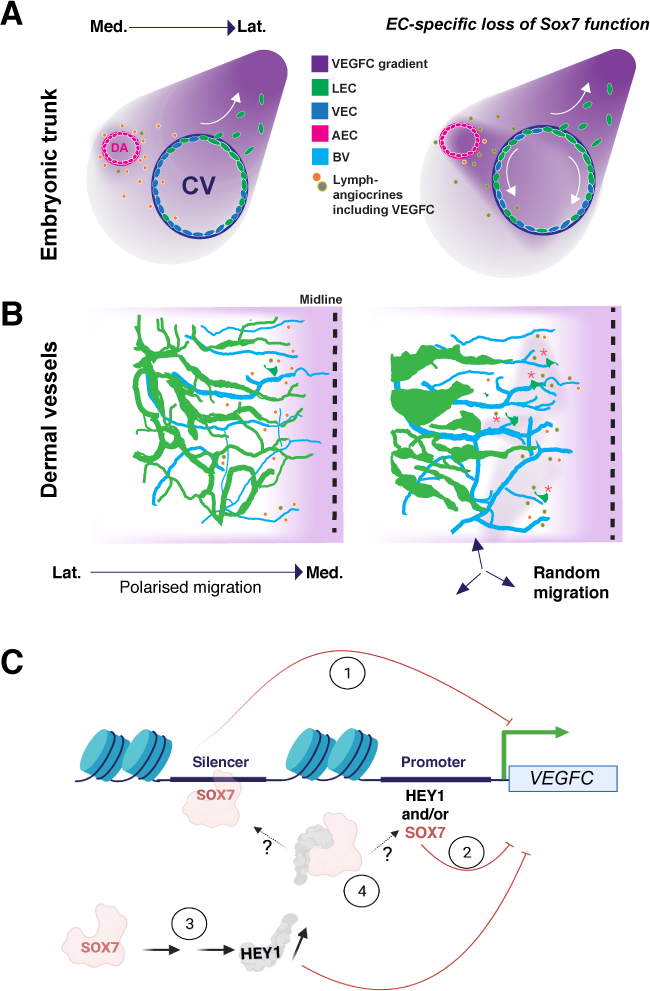
Model depicting how loss of SOX7 function in blood vascular endothelial cells causes defective lymphatic vascular patterning. (A and B) LEC progenitor organisation is impaired in both early stage and organ-specific lymphangiogenesis. (**A**) In physiological conditions, LEC progenitors emerge in the dorsal lateral part of the CVs and migrate towards the dorso-lateral aspects of the embryo. In the absence of functional SOX7, blood vascular endothelial cells from the arterial compartment increase the local expression of endothelial VEGFC, perturbing its tissue distribution. This induces and/or expands the number of LEC progenitors in ventro-medial aspects of the CVs. (**B**) Dermal lymphatics are dysmorphic in the absence of SOX7 function in blood vascular endothelial cells. An increase in VEGFC in the dermal blood endothelial cells causes hyperproliferation of local LEC progenitors. This is shown by an increase in the emergence of local LEC progenitors from the endomucin-positive blood capillary plexus at E12.5 (red asterisks). Changes in expressions of other SOX7-dependent lymphangiocrines also contribute to the migration defects in the dermal lymphatics. (**C**) Model showing the molecular mechanisms of SOX7-dependent repression of *VEGFC* transcription identified in this study: 1) SOX7 binds to distal regulatory elements (silencing regions) to suppress *VEGFC* transcription. 2) SOX7 represses VEGFC promoter activity. 3) SOX7 acts upstream of Notch effector and repressor, HEY1. SOX7-induced expression of HEY1 causes *VEGFC* downregulation and 4) Protein-protein interaction between SOX7 and HEY1 forms a complex to repress *VEGFC* transcription. LEC, lymphatic endothelial cells; AEC, arterial endothelial cells; VEC, venous endothelial cells; Lat., lateral; Med, medial.

To determine if the putative regulatory elements observed in the human ChIP-Seq dataset are conserved in mouse *in vivo*, we used the *Sox7*-V5 reporter line to perform ChIP-Seq analysis in E9.5 and E10.5 mouse embryos. This approach identified 12976 binding regions in E9.5 and 3351 binding regions in E10.5 embryos (Sup. Fig. 6B). In addition to observing the presence of SOX7 binding motifs amongst the most enriched transcription binding motifs at both time points (Sup. Fig. 6C–E), we further validated this dataset by performing ChIP-PCR on the top 5 most intense peaks and other assumed unbound regions (negative control) (Sup. Fig. 6F). Gene ontology analysis found GO:0007411 axon guidance and GO:0045499 chemical repellent activities as the top biological process and molecular function in the overlapping genes between E9.5 and E10.5 SOX7-V5 ChIP-Seq (Sup. Fig. 6 G-J). Pathway analysis further showed that “axon guidance” is the most significant pathway associated to these genes, suggesting that SOX7 may also regulate lymphatic patterning through the modulation of other guidance molecules in addition to VEGFC (Sup. Table 1).

A common feature of the *in vivo* E9.5 and E10.5 ChIP-Seq results and the SOX7 HUVEC ChIP-Seq dataset is the presence of peaks assigned to the *Vegfc* locus. In contrast, we did not detect any binding to regions associated to *Vegf*, *Vegfd*, *Vegfr1*, *Vegfr2* or *Vegfr3*. At E9.5, SOX7 binds to sites 186 kb upstream (*Vegfc*-186) and 251 kb downstream (*Vegfc*+251) of the VEGFC TSS. At E10.5, SOX7 binds to sites 381 kb and 152 kb upstream (*Vegfc*-381, *Vegfc*-152) of the VEGFC TSS, suggesting 4 putative regulatory elements are recruited differentially at distinct embryonic time points. Of these 4 binding regions, *Vegfc*-152 is associated with a region enriched for the repressive histone mark H3K9me3, typically linked to a constitutive heterochromatin state as revealed by the E11 mouse ENCODE data repository (Fig. 4D). Other peaks do not appear to coincide with any of the known histone marks reported in the ENCODE database for E10.5-E11 mouse embryos.

Both *VEGFC-255* (human) and *Vegfc-152* (mouse) contain multiple partially conserved SOX and ETS binding motifs (Sup. Fig. 7). Identification of SOX7 binding sites in distal regions assigned to *VEGFC* loci, consistently in both human and mouse, suggests that SOX7 might act as a direct transcriptional regulator of VEGFC in both species through these putative distal regulatory elements.

To test whether *VEGFC-255* and *Vegfc-152* have any regulatory potential on VEGFC promoter activity, we generated a series of VEGFC luciferase reporter constructs that harbour a combination of different putative regulatory regions from mouse (mP) and Human (hP) (Fig. 4C and 4E, Sup. Fig. 7). As a starting point, we used SOX7-chromatin bound regions “B” (black bar, Fig 4C, E and F) in combination with a conserved flanking region “C” (blue bar, Fig. 4C, 4E and 4F) located in 3’ of the “B” region in both human and mouse. In addition, we also included in these reporter constructs nearby genomic regions labelled by repressive marks referred to as “R” (pink bar, Fig. 4C, E and F). The “R” region is located in 5’ of the B element in mouse and in 3’ in human. The largest fragment generated for this series of combinations is made up of all “RBC” regions and is approximately 6.5 kb long.

As noted above, SOX7 repressed both human and mouse VEGFC basal promoter activity (Fig. 4A, F). For the human constructs addition of the “R” and “B” and “C” regions in all cases led to a decrease of VEGFC basal promoter activity. This repressive effect was further enhanced in presence of SOX7 over-expression (Fig. 4F). By contrast in the context of the mouse genome, the “RBC” region caused an increase of VEGFC basal promoter activity, however SOX7 over-expression led to a marked decrease of the VEGFC reporter activity, similar to what was observed with the human constructs. Taken together, results from this *in vitro* reporter assay support the notion that SOX7 directly represses *Vegfc* transcription via a combination of inhibitory regions, likely involving the coordinated activity of both the VEGFC promoter as well as distant regulatory elements. This repressive mechanism appears to be functionally conserved between mouse and human.

More broadly, the observation that SOX7 loss-of-function leads to *Vegfc* upregulation both *in vitro* and *in vivo*, combined with the identification of SOX7-dependent VEGFC silencing regions, suggests that direct transcriptional repression by SOX7 is a significant mechanism of VEGFC regulation in BECs.

### Loss of *Sox7* function disturbs the emergence and expansion of LEC progenitors *in vivo* similar to VEGFC gain-of-function

Having already established that BEC-specific loss of SOX7 function in mice causes defects in lymphatic patterning, we next sought to link these defects specifically with VEGFC de-repression. Previous work has established that VEGFC promotes LEC specification and expansion (Mäkinen et al., 2001; Pichol-Thievend et al., 2018; Srinivasan et al., 2014). We therefore hypothesised that ectopic activation of VEGFC, especially in close vicinity to the CVs at the time of the initial steps of lymphangiogenesis, would change both the distribution and number of PROX1-positive LEC progenitors. To test this hypothesis, we analysed both the anatomical locations and density of LEC progenitors in the CVs in E11.5 *Sox7^iECKO^* mutant embryos. During development, PROX1+ LEC progenitors are polarised to a dorsal-lateral aspect of the CVs in the anterior part of the embryo (Bowles et al., 2014; Wigle and Oliver, 1999). Our analysis revealed an increase in the proportion of PROX1+ LEC progenitors in the *Sox7^iECKO^*, as well as an ectopic distribution of LEC progenitors in the ventro-medial side of the CVs (yellow arrowheads, Fig. 5A-C, Sup. Fig. 4H).

Previously, we and others have shown that the dermal blood capillary network is an on-going source of the dermal LECs progenitors and emergence of these precursor cells from the endomucin-positive plexus is dependent on CCBE1/VEGFC/VEGFR3 signalling (Martinez-Corral et al., 2015; Pichol-Thievend et al., 2018). High levels of VEGFC expression in blood vascular endothelial cells increases the number of LECs within the blood capillary plexus, as observed in the *Tie2Cre;VegfcGOF* mice (Pichol-Thievend et al., 2018). To investigate whether an increase of *Vegfc* levels in *Sox7^iECKO^* mutants (as characterised above in Fig. 2C) similarly affects the number of LECs in the dermal capillary, we quantified the number of isolated PROX1-positive LEC progenitors. These progenitors were located either on or within the vessel wall of endomucin-positive blood vessels in E12.5 skin (Fig. 5D, E, Sup. Fig. 8). Our quantitative analysis focussed on EMCN^low/-^PROX1^+^NRP2^+/low^ individual LECs (arrowhead, type I) on blood vessels, EMCN^+^PROX1^+^ NRP2^low/-^ LECs within the vessel wall (arrow, type II) and EMCN^+/-^ PROX1^+^NRP2^+/low^ LEC clusters (2-4 cells, empty arrowhead, type III) that were exiting from the blood plexus network. Similar to the *Tie2Cre;VegfcGOF* embryonic skin, analysis of *Sox7^iECKO^* skin revealed a significant increase in the number of PROX1+ LEC progenitors associated with the blood plexus (Fig. 5E, Sup. Fig. 8C-E).

Taken together, the increase of LEC specification events in the CVs and the dermal blood capillaries correlated with higher levels of *Vegfc* transcripts in SOX7 mutants *in vivo*, establishes SOX7 as a key BEC-specific regulator of lymphatic vessel patterning through negative modulation of VEGFC expression.

## Discussion

While it is well established that SOX7 acts in a cell-autonomous manner to instruct angiogenesis and arteriovenous specification during development (Hermkens et al., 2015; Kim et al., 2016; Lilly et al., 2017), this study reveals for the first time a function for this BEC-specific transcription factor in regulating signalling crucial for the organisation of the lymphatic vasculature. We show that one mechanism by which SOX7 achieves this role is via the transcriptional repression of *Vegfc* in the blood vascular endothelium. This fine-tunning of VEGFC local repression is required to maintain strict control over the number and spatial distribution of LECs essential for correct assembly of the dermal lymphatic vascular network.

To date, only a handful of factors have been shown to transcriptionally regulate VEGFC expression (Cohen et al., 2020; Gauvrit et al., 2018; Schuermann et al., 2015), with no previous study describing an endothelial-specific and direct molecular mechanism that regulates VEGFC transcription. Our work sheds light into how the BEC-specific SOX7 transcription factor dampens local VEGFC levels to modulate tissue growth in a paracrine manner likely via a combination of mechanisms: 1) indirectly through recruitment and transcriptional control of the repressor protein HEY1 2) directly via binding to silencing elements associated with the *Vegfc* locus.

Conventionally, within the context of embryonic development, the SOXF (7, -17 and 18) transcription factors have been shown to act mainly as transcriptional activators (Chew and Gallo, 2009). Unlike the SOXB2 transcription factors (SOX21 and SOX14) that possess a repression domain, SOXF proteins have a transactivating domain (Schock and LaBonne, 2020; Uchikawa et al., 1999). Similar to SOX18 and SOX17 (Corada et al., 2013; François et al., 2008; Hosking et al., 2004), SOX7 has primarily been reported to positively regulate gene transcription (Chiang et al., 2017; Costa et al., 2012; Futaki et al., 2004; Kim et al., 2016; Murakami et al., 2004; Niimi et al., 2004; Sacilotto et al., 2013), although there have been a handful of reports of SOX7 acting to repress transcription via the sequestration of activators through direct protein-protein interaction (Fan et al., 2018; Guo et al., 2008; Lilly et al., 2016). Other repressive mechanisms are possibly at play such as an indirect regulatory cascade whereby SOX7 loss-of-function causes a failure to activate a repressor such as HEY1, which in turn causes an activation of downstream target genes. It is also possible that the DNA environment and the code of binding cofactors turn an activator into a repressor, as exemplified by a Drosophila morphogen, Dorsal (Ip, 1995). Here we show that repression of *Vegfc* transcription by SOX7 may involve indirect mechanisms mediated by the modulation of Notch signalling effector activity, but importantly we also uncover a novel mechanism involving direct recruitment of SOX7 to silencing elements associated with the *Vegfc* locus.

In terms of indirect regulation of VEGFC expression by SOX7, while this study has not definitively defined the role of SOX7-HEY1 interaction in the context of *VEGFC* transcriptional regulation, our data provide compelling evidence to explore this interaction further. It is possible that SOX7 may induce HEY1 gene expression to promote the assembly of a SOX7/HEY1 heterodimer that in turn represses VEGFC as part of a positive feedback loop. Notably, we detected two HEY1 binding motifs 5’-CACGTG-3’ in human *VEGFC-255* RBC and a single HEY1 binding motif in the human VEGFC promoter (data not shown). Although the SOXF and HEY1 binding sites are far apart, physical interaction could be made possible through changes in chromatin conformation. In addition, it is possible that SOX7 recruitment of its protein partner does not involve direct HEY1/DNA interaction. Importantly, this study supports a previous report of the importance of Notch signalling in the negative regulation of LEC specification and expansion (Murtomaki et al., 2013), and begins to unravel the potential role of SOX7 in mediating this regulation. We propose that SOX7 and Notch/HEY1 may function as converging pathways, to produce a combinatorial effect on *Vegfc* repression in order to maintain LEC progenitor homeostasis.

In parallel to indirect regulation of *Vegfc* by SOX7, our study identified putative VEGFC silencing elements in both the human and mouse genomes, and suggest that SOX7 can regulate *Vegfc* transcription through direct binding at these sites. While a handful of factors have been shown to transcriptionally regulate VEGFC expression in the past (Cohen et al., 2020; Gauvrit et al., 2018; Schuermann et al., 2015), this is the first identification of an endothelial-specific and direct molecular mechanism able to regulate *Vegfc* transcription.

As previously described, silencing elements are enriched in the H3K9me3 mark, and H3K27me3 mark which is associated with Polycomb repression (Pang and Snyder, 2020). Deletion of Polycomb-bound elements was recently shown to lead to de-repression of their target genes, inferring that Polycomb-bound regions could potentially function as silencers (Ngan et al., 2020). Having identified a number of SOX7 binding sites associated with the VEGFC genomic locus, *in silico* evaluation of the chromatin states of the *VEGFC-255* and *Vegfc-152* regions revealed characteristics of silencing elements. We further substantiated these *in silico* observations by demonstrating the ability of these genomic regions to reduce VEGFC promoter activity *in vitro* when fused to luciferase synthetic constructs. The presence of insulator CTCF and Polycomb repressor binding sites in *VEGFC-255* suggest a possible direct or indirect molecular relationship between SOX7 and CTCF or Polycomb repressing complexes to elicit repression. Interestingly, both CTCF and CTCFL (BORIS) are amongst the top enriched secondary binding motifs in SOX7 HUVEC ChIP-Seq (Sup. Fig. 9A). Further, overlapping of the SOX7 HUVEC and V5 ChIP-Seq data with histone marks from the ENCODE consortium reveals a large proportion of the peaks have at least 50% overlap with a repressive mark (H3K27me3 or H3K36me3) further supporting a repressive mode of action for SOX7 (Sup. Fig. 9B-L). Analysis of the chromatin state of the SOX7 HUVEC ChIP-Seq reveals most peaks are associated to heterochromatin, followed by strong enhancers and weakly transcribed regions (Sup. Fig. 9F). Interestingly, 4.7% of the peaks are found associated to Polycomb-repressed regions.

A repressive function has been implicated for at least one other SOXF transcription factor; with SOX18 homodimer binding motifs (inverted repeat-5, IR-5) found associated with both active and repressive histone marks (Moustaqil et al., 2018). SOX7 repressive effects on *Vegfc* transcription in the CVs and dermal blood plexus could potentially act as a “shield” to maintain some LEC-free areas, therefore preserving vessel integrity and function (see model, Fig. 6). Nonetheless, it is unlikely that the SOX7 lymphatic defects is limited to just an upregulation of VEGFC level. Identification of multiple putative guidance cues previously implicated in LEC migration and patterning such as semaphorins and netrins in SOX7-V5 ChIP-Seq data further suggested that dysregulation of these guidance cues could also contribute to the overall migration defects observed in the lymphatic network (Liu et al., 2016; Uchida et al., 2015).

Whether SOX7 functions as an activator or a repressor in a given context, is likely to be dictated by the surrounding chromatin state as well as the spatial and temporal distribution of its binding co-factors (Ip, 1995). Like the CRX transcription factor in rod photoreceptors, the SOX7 dual-function as activator or repressor might also be dependent on the number and affinity of its binding sites at particular regulatory elements (White et al., 2016). Given the known role of SOX7 to maintain endothelial cell identity and repress the hemogenic program (Gandillet et al., 2009; Lilly et al., 2016), it is highly likely that the coordinated activity of activating and repressive function are at play in the same cell at the same time for a specific subset of gene regulatory network.

By uncovering a number of novel mechanisms by which SOX7 can modulate *Vegfc* transcription, this study reveals that the final transcriptional output of VEGFC in BECs is likely to involve a complex interplay of molecular events exquisitely coordinated in a strict spatio-temporal manner. Our data suggest that while SOX7 alone appears sufficient to reduce VEGFC promoter activity, it is likely that *in vivo* SOX7 works in concert with other cofactors at different distal regulatory elements to effectively coordinate changes in *Vegfc* transcription rate. We demonstrate that this BEC-specific SOX7/VEGF-dependent regulatory axis, potentially with other unidentified SOX7 regulated lymphangiocrines, are essential for appropriate LEC specification and spatial organisation into a functional lymphatic network; further cementing the role of the blood vascular network in guiding lymphangiogenesis.

## MATERIALS AND METHODS

### Mice and breeding

All animal work conformed to ethical guidelines and was approved by the relevant University of Queensland and University of Sydney ethics committee. The following mice used in this study have been reported previously: *Sox7:tm1* (Chiang et al., 2017), *Cdh5-CreERT2* (Wang et al., 2010), *Sox7^fl/fl^* (Lilly et al., 2017) and *mTmG* (Schimmel et al., 2020). The *Sox7-V5* tagged mice (mixed background) were generated as detailed in Supplemental Materials and Methods. The *Sox7* endothelial-specific conditional knockout, *Cdh5-CreERT2:Sox7 ^fl/fl^*, was a crossed between *Cdh5-CreERT2* and *Sox7^fl/fl^*. All mice strains were in C57BL/6 genetic background.

To delete *Sox7* specifically in endothelial cells, each pregnant dam received two consecutive intraperitoneal injections of tamoxifen (T5648, Sigma Aldrich), at 1.5 mg per pulse at the indicated time point. Tamoxifen was reconstituted in 10% (v/v) ethanol and 90% (v/v) peanut oil.

### Chromatin immunoprecipitation (ChIP) sequencing

Homozygous *Sox7*-V5 mouse embryos confirmed by sequencing and PCR genotyping were collected at E9.5 and E10.5, and processed as detailed in Supplemental Materials and Methods (Mohammed et al., 2013). DNA amplification was performed using TruSeq ChIPseq kit (Illumina, IP-202-1012), following an immunoprecipitation step using 10 μg V5 monoclonal antibody (R960CUS, Invitrogen). The library was quantified using the KAPA library quantification kit for Illumina sequencing platforms (KAPA Biosystems, KK4824) and 50bp single end reads were sequenced on a HiSeq2500 following the manufacturer’s protocol. FASTQ files were mapped to mouse the reference genome Genome Reference Consortium (GRC)m38/mm10 (released in Dec 2011) using Bowtie2 (Galaxy version 2.2.6.2) (Langmead and Salzberg, 2012). Peaks were called using MACS version 2.1.0 (Zhang et al., 2008). Potential binding regions of SOX7 (or peaks) were called using each corresponding input DNA as a reference, with a false discovery rate (FDR) of 0.01. Sequencing was performed by the IMB Sequencing Facility, University of Queensland. Please refer to Supplemental materials and Methods for more details.

### Tissue sectioning and immunofluorescence staining

For tissue sectioning, whole embryo analysis and whole-mount immunostaining of embryonic skin, whole embryos were fixed in 4% (w/v) paraformaldehyde/PBS overnight at 4°C, and washed three times in PBS. Whole-mount immunofluorescence staining of embryonic skins was performed as described (Pichol-Thievend et al., 2018). For sectional analysis, embryos were embedded in 4% agarose and sectioned transversely at 150μm using the Leica VT1000 S vibrating microtome. For paraffin sectioning, fixed embryos were dehydrated and cleared with a tissue processor (Leica Biosystem). Embryos were then embedded in paraffin before sectioned (8 μm) with a microtome (Polanco et al., 2010). Sections were next rehydrated, boiled in appropriate antigen retrieval solution (Vector) before blocking with blocking solution. This was followed by primary and secondary antibody incubation as described (Pichol-Thievend et al., 2018). Details about antibodies can be found in supplemental Materials and Methods.

### Co-immunoprecipitation and western blotting

At 48h post-transfection, HEK293 cells were rinsed with PBS before lysis and protein extraction with NE-PER Nuclear and Cytoplasmic Extraction Reagents (Pierce). Protein concentration was measured with Bicinchoninic Acid (BCA) assay (Pierce). To prevent protein degradation, cOmplete Protease Inhibitor Cocktail (Roche, 11697498001) was added to the cell lysate. Clarified nuclear fractions containing the HEY1 and SOX7 were then incubated with anti-GFP (3E6) (1:1000, A-11120, Thermo-Scientific) or random IgG bound magnetic Dynabeads for 2-3 h, at 4°C with rotation. Beads were then washed 3 times in RIPA lysis buffer on a magnetic stand, then eluted in SDS sampling buffer with heating for 5 min, before analysis by 12% SDS-PAGE. Immunoblotting was performed with chemiluminescent HRP detection (Pierce) using a ChemiDoc^TM^ Imaging Systems (Bio-Rad). Pull-down and immunoblotting assays were modified from (Schmidt et al., 2009). Details about antibodies can be found in supplemental Materials and Methods.

### Fluorescent Activated Cell sorting (FACS) and expression analysis

CD31+CD45- endothelial cells were isolated from skin flaps of E14.5 sibling controls and *Sox7^iECKO^* mutants (treated with tamoxifen at E11.5, E12.5) (Kazenwadel et al., 2012). RNA was extracted, amplified and cDNA was synthesised as previously described (Chiang et al., 2017). Primer sequences and details of the qPCR analysis are provided in the supplemental Materials and Methods.

### Cell culture and siRNA mediated silencing experiment

Human umbilical arterial endothelial cells (HUAECs) (C-12202, PromoCell) and human umbilical venous endothelial cells (HUVECs) (CC-2519A) were maintained in EGM2 media (Lonza). For siRNA, Smartpool: ON-TARGETplus Human SOX7 (83595) siRNA 50 nM (containing a mixture of 4 siRNAs against *SOX7*) (Dharmacon) and scrambled siRNA (Dharmacon) were used. HUVECs and HUAECs were transfected at ∼40-50% confluency, using Lipofectamine^TM^ 3000 (Invitrogen) in OptiMEM media. Cells were harvested for RNA extraction at 30 h or 17 h post-transfection.

HEK293 cells were maintained in DMEM added with 10% FCS and 1% Penicillin/Streptomycin. HEK293 cells for co-immunoprecipitation assays were transfected for 6 h and incubated for another 18 h before harvesting. DNA transfection was performed using X-tremeGENE^TM^ 9 DNA transfection reagent (Roche) in OptiMEM media according to the manufacturer’s instructions.

### Imaging and data analysis

Images were captured using a Zeiss LSM710 META BIG, Zeiss LSM 710 FCS or Leica TCS SP8 HyD confocal microscopes with the 10X, 20X or 40X oil objectives. Images were analysed with Zeiss Zen software, the Bitplane IMARIS suite and Image J/FIJI. All graphs and statistical tests were performed with GraphPad Prism 8 and illustrated with Adobe Illustrator CS6. Details about quantification can be found in Supplemental Materials and Methods. Imaging was performed in the Australian Cancer Research Foundation (ACRF)’s Dynamic Imaging Facility at the Institute for Molecular Bioscience (University of Queensland) and the Sydney Cytometry facilities at Centenary Institute (University of Sydney).

## Supplemental Materials and Methods

### Mice and generation of *Sox7*-V5

All animal work conformed to ethical guidelines and was approved by the relevant University of Queensland ethics committee.

The *Sox7:tm1* embryonic stem cells were generated using BAC recombination, and then injected into C57BL/6 blastocysts (Knockout Mouse Project (KOMP) repository, US). The *Sox7:tm1* has the entire *Sox7* HMG domain in exon 1 and part of the exon 2 displaced by a *lacZ* cassette. The resulting *Sox7^+/-^* carriers were crossed to generate *Sox7^-/-^* embryos.

To identify *Sox7^fl^* progeny of *Cdh5-CreERT2:Sox7 fl/fl* line, genotyping primers were:

m*Sox7fl*_F: 5’-GGGTTACCGCACTTAAGAGACA-3’

m*Sox7fl*_R: 5’-GGAAGTCCTACCCGACCTAATC-3’

Wild-type band was 195 bp, flox band was 345 bp.

To identify *Cdh5-CreERT2* progeny, genotyping primers were:

*Cre_F*: 5’-CTGACCGTACACCAAAATTTGCCTG-3’

*Cre_R*: 5’-GATAATCGCGAACATCTTCAGGTTC-3’

Cre positive band was 200 bp.

*Sox7*-V5 tagged mice were generated by CRISPR according to the published protocol (Yang et al., 2013) using the following primer and donor oligo:

*Sox7*-CRISPRgRNA1: 5’-GCTACAGTGTGTCATAGAGC-3’

*Sox7*-V5 donor oligo (V5 tag = underlined):

5’-AGCCTCATCTCAGTCCTGGCTGATGCCACGGCCACGTATTACAACAGCT ACAGTGTGTCAGGCAAGCCCATCCCCAACCCCCTGCTGGGCCTGGACAGCACCTAGAG CTGGAGGAATGGAGCCTGGCCCAGCCCTGCCATCCCCTCCTCCCTATGAAGCACT-3’ CBB6F1/BCB6F1 females (CBA x C57BL/6) and Arc:Arc(S) females were used as embryo donors and foster mothers, respectively. Super-ovulated CBB6F1/BCB6F1 females were mated to CBB6F1/BCB6F1 studs, and fertilised embryos were collected subsequently from the oviducts. CRISPR injection-mix containing (10 ng/ul of Cas9 mRNA, 5 ng/ul of gRNA1 and 20 ng/ul of donor oligo) was injected into the pro-nuclei of fertilised eggs. The injected zygotes were cultured in KSOM/M16 media at 37°C, 5% CO_2_ until a two-cell stage. Subsequently, 15-25 two-cell zygotes were transferred into the uterus of pseudo pregnant Arc:Arc(S) at 0.5 dpc.

Genotyping primers for *Sox7*-V5 were:

*Sox7*-V5_SF: 5’-TCACACCTAGTCCCCTCCAC-3’

*Sox7*-V5_SR: 5’-GTGGGGTTTTGCCAGTTAGA-3’

*Sox7*-V5_V5R: 5’-TCCAGCTCTAGGTGCTGTCC-3’

For SF/SR primer pairs:

The wild type allele corresponds to a band at 820 bp, V5 band was 862 bp.

SF/V5R primer pairs

V5 positive band was 479 bp

### Nuclei isolation for snRNA-Seq

Skins from E14.5 wild-type mouse embryos were dissected, and snapped freeze in liquid nitrogen before stored in -80°C freezer. Skins from 3 separate embryos were pooled, and single nuclei were extracted and isolated. Briefly, skins were homogenised in chilled nuclei EZ lysis buffer (N3408, Sigma Aldrich) using a 1 ml douncer, suspended with a wide bore tip before passing through a 70 μm cell strainer (352350, Falcon). Nuclei were washed and resuspended in PBS buffer containing 1% BSA, 0.2 U/μl RNAse inhibitor (10777019, Invitrogen), stained in 10 μg/mL DAPI, and passed through a 40 μm cell strainer (352340, Falcon). Nuclei were then sorted using BD FACSAria III with 70 μM nozzle, before loaded onto the 10X Chromium single cell chip (v3, 10X Genomics).

### Bioinformatic analysis of single-nuclei RNA-Seq

Single nuclei were loaded onto 10× Chromium platform for single nucleus gene expression assay and the snRNA-seq library was sequenced on Illumina NovaSeq 6000. The raw sequencing data was processed using Cell Ranger (version 3.1.0) (Zheng et al., 2017), which takes care of read quality control (QC), demultiplexing, genome alignment and quantification. The resulting count matrix with the dimension number of barcodes × number of transcripts was obtained and loaded in R (version 4.0.4) for downstream analysis using Seurat (version 4.0.3) (Butler et al., 2018), including the selection and filtration of nuclei based on QC metrics, normalization and variance stabilisation of molecular count data using sctransform, dimensionality reduction by Principal Component Analysis and Uniform Manifold Approximation and Projection embedding, and clustering analysis.

In the step of QC, nuclei expressing <200 genes or with genes that were expressed in <3 cells were filtered. In addition, nuclei were further assessed based on the following criteria: the ones that have >5% mitochondrial counts and expressed <200 or >3,500 unique genes were excluded. This results in 8,380 quality cells. In the step of clustering analysis, the top 30 principal components were used as input. At a resolution of 0.8, 26 clusters were retrieved and projected onto UMAP plots. Following clustering analysis, genes differentially expressed in each of the clusters were determined using a method of differential expression analysis based on the non-parametric Wilcoxon rank sum test, and genes were filtered based on a minimum log2 foldchange (0.25) for average expression of gene in a cluster relative to the average expression in all other clusters combined. The top 30 genes were used for cell type annotation. Visualisation of expression patterns of specific genes across clusters was performed with Nebulosa in Seurat (Alquicira-Hernandez and Powell, 2021).

### Microarray samples

Embryonic mouse LECs and BECs were purified from E14.5, E16.5 and E18.5 skin as previously described (Kazenwadel et al., Blood, 2010). Each sample was generated from 15-21 embryos (pooled from multiple litters) and three independent samples were generated for each timepoint. Between 0.5 and 3 ug of total RNA was isolated per sample using TRIzol reagent (Thermo Fisher), according to the manufacturer’s directions. RNA quality was assessed using a Bioanalyser (Agilent Technologies); all samples achieved an RNA Integrity Number (RIN) score > 9.5. Samples were submitted to the ACRF Cancer Genomics Facility (Adelaide, Australia) and hybridised to GeneChip® Mouse Gene 1.0 ST arrays (Affymetrix) for gene expression profiling. Microarray data analysis was performed using Partek Genomics Suite™ version 6.4 software (Partek Incorporated, St. Louis, MO). Differential gene expression was assessed by one-way ANOVA, with *P*-values adjusted for multiple testing using a step-up false discovery rate correction (Benjamini and Hochberg, 1995).

### Fluorescent RNA *in situ* hybridisation

Embryonic tissue sections cut at 30 µm thickness were processed for fluorescent RNA *in situ* hybridisation using the commercially available RNAscope^®^ Multiplex Fluorescent v2 assays (Advanced Cell Diagnostics (ACD)) according to the manufacturer’s protocols for fresh frozen samples. For detection of *Vegfc* RNA expression the Mm-Vegfc-C2 probe (Cat No. 492701-C2; ACD) was used. Detection was performed using the Opal™ 620 dye (FP1495001KT, Perkin Elmer) at a final concentration of 1:750 in TSA buffer provided in the RNAscope® Multiplex Fluorescent v2 kit.

For subsequent immunostaining the sections were rehydrated in PBS, then permeabilised in 0.5% Triton X-100 in PBS (PBS-T) followed by blocking in PermBlock (3% BSA, 0.5%Tween-20 in PBS) and incubation with primary antibody diluted in PermBlock at 4°C. Following 3 washing steps with PBS- T, tissues were incubated in secondary antibodies labelled with Alexa- dyes (Life Technologies) at 4°C. After 3 washing steps with PBS- T, the sections were mounted with Mowiol. The following antibodies were used: goat polyclonal anti- human PROX1 (AF2727, R&D Systems), rat monoclonal anti- mouse PECAM- 1 [clone 5D2.6 and clone 1G5.1 (Wegmann et al., 2006) and rabbit anti-GFP (ab6556, Abcam).

### Generation of expression and luciferase constructs

#### Expression constructs

To generate the expression constructs for co-immunoprecipitation assays, the mouse SOX7 and HEY1 full-length regions were amplified from a human cDNA library generated in-house from HUVECs. To generate GFP-SOX7, we used the following primers (restriction sites were underlined):

GFP-SOX7_F (HindIII): 5’-CGTA<u>AAGCTT</u>CGATGGCTTCGCTGCTGGG-3’

GFP-SOX7_R (BamH1):

5’-GATC<u>GGATCC</u>CTATGACACACTGTAGCTGTTGTAGT-3’

To generate SOX7-GFP, primers used were:

SOX7-GFP_F (HindIII): 5’-CGTA<u>AAGCTT</u>ATGGCTTCGCTGCTGGGAGC-3’

SOX7-GFP_R (BamHI):

5’-GATC<u>GGATCC</u>CGTGACACACTGTAGCTGTTGTAGT-3’

To generate Cherry-HEY1, primers used were:

Cherry-HEY1_F (HindIII): 5’-CGTA<u>AAGCTT</u>CGATGAAGCGAGCTCACCCC-3’

Cherry-HEY1_R (BamHI): 5’-GATC<u>GGATCC</u>AAAAGCTCCGATCTCCGTCC-3’

To generate HEY1-Cherry, primers used were:

HEY1-Cherry_F (Xho1): 5’-GATC<u>CTCGAG</u>ATGAAGCGAGCTCACCCC-3’

HEY1-Cherry_R (HindIII): 5’-CGTA<u>AAGCTT</u>AAAAGCTCCGATCTCCGTCC-3’

Full-length SOX7 was sub-cloned into linearised pEGFP-N1 and pEGFP-C1 (Clontech), respectively to generate SOX7-GFP and GFP-SOX7. Full-length HEY1 was sub-cloned into linearised pmCherry-N1 and pmCherry-C1 (Clontech) to generate HEY1-Cherry and Cherry-HEY1.

To generate expression constructs for luciferase assay,

Full-length HEY1 was amplified from human cDNA library with primers:

pCDNA HEY1_F (EcoRI): 5’-CGTA<u>GAATTC</u>ACCATGAAGCGAGCTCACCCC-3’

pCDNA HEY1_R (XhoI):

R: 5’-GATC<u>CTCGAG</u>TTAAAAAGCTCCGATCTCCGTCC-3’

And sub-cloned into linearised pCDNA3.1 (Invitrogen) to generate pCDNA HEY1.

Full-length SOX7 was amplified from human cDNA library with primers:

pCDNA SOX7_F (HindIII):

5’-CGTA<u>AAGCTT</u>GCC ATGGCTTCGCTGCTGGGAGC-3’

pCDNA SOX7_R (BamHI):

5’-GATC<u>GGATCC</u> CTATGACACACTGTAGCTGTTGTAGTACGT-3’

And sub-cloned into linearised pCDNA3.1glomyc to generate pCDNA SOX7.

#### Luciferase constructs

Human VEGFC promoter (hP) was the pGL3-VEGFC promoter-luc-3.4kbHindIII from Kari Alitalo (Chilov et al., 1997). Mouse *Vegfc* promoter (mP) was pGL4-VEGFC promoter-luc-370 from Min-Jen Hsu (Huang et al., 2014), sub-cloned into linearised pGL3-basic using primers containing HindIII and XhoI restriction enzyme sites (underlined):

–370 mP_Vegfc_F(XhoI): CTAGCCCGGG<u>CTCGAG</u>TCTTCGGGACGAGTGGAAC

+1 mP_Vegfc_R(HindIII): CCGGAATGCC<u>AAGCTT</u>TGGTGGATGGACCGGGAG

To generate mouse *Vegfc-152* BC-mP-luc construct, the 3.577 kb mouse *Vegfc-152* BC region (GRCm38/mm10_dna range: Chr8:53,925,003-53,928,579) and 371 bp mouse VEGFC promoter (-370/+1) was subcloned into the linearised pGL3-basic, with primers (restriction sites were underlined):

#### Insert 1 (*Vegfc-152* BC)

*mVegfc-152* BC-mP-luc_1F (XhoI):

5’-GCGTGCTAGCCCGGG<u>CTCGAG</u>TGATGACTAGTGTTCAATTCCTTT-3’

*mVegfc-152* BC/RBC-mP-luc_1R:

5’-GTCCCGAAGACATAGATATTGAGATGCCATCTTTT-3’

#### Insert 2 (VEGFC promoter -370/+1)

m*Sox*BC/RBC(*Vegfc-*370/+1)_2F:

5’-AATATCTATGTCTTCGGGACGAGTGGAAC-3’

m*Sox*BC/RBC(*Vegfc-*370/+1)_2R (HindIII):

5’-CAGTACCGGAATGCC<u>AAGCTT</u>GGTGGATGGACCGGGAG-3’

To generate mouse *Vegfc-152* RBC-mP-luc construct, the 6.244 kb mouse *Vegfc-152* BC region (GRCm38/mm10_dna range: Chr8:53,922,336-53,928,579) and 371 bp mouse VEGFC promoter (-370/+1) was subcloned into the linearised pGL3-basic, with primers (restriction enzyme sites underlined):

#### Insert 1 (*Vegfc-152* RBC)

*mVegfc-152* RBC-mP-luc_1F:

5’-GCGTGCTAGCCCGGGCTCGACAGAACAGGTAGCAGAGGCTA-3’

*mVegfc-152* BC/RBC-mP-luc_1R:

5’-GTCCCGAAGACATAGATATTGAGATGCCATCTTTT-3’

#### Insert 2 (VEGFC promoter -370/+1)

m*Sox*BC/RBC(*Vegfc-*370/+1)_2F:

5’-AATATCTATGTCTTCGGGACGAGTGGAAC-3’

m*Sox*BC/RBC(*Vegfc-*370/+1)_2R:

5’-CAGTACCGGAATGCCAAGCTTGGTGGATGGACCGGGAG-3’

pGL3-basic was linearised with XhoI-HF and HindIII-HF.

To generate human *VEGFC-255* BC-hP-luc construct, the 2.014 kb human *VEGFC-255* BC region (GRCh37/hg19_dna range: chr4:177,968,343-177,970,356) was subcloned into the Kpn1 linearised pGL3-VEGFC promoter-luc-3.4kbHindIII plasmid, with primers (restriction sites were underlined):

*hVEGFC-255* BC/RBC-hP-luc_F (KpnI):

5’-TTTCTCTATCGATA<u>GGTACC</u>AAAGGGTCAAATCGCACAATGC-3’

*hVEGFC-255* BC-hP-luc_R (KpnI):

5’-TCGCGTAAGAGCTC<u>GGTACC</u>CAATTGTGCAAGAAGGAGCTTGAG-3’

To generate human *VEGFC-255* RBC-hP-luc construct, the 6.503 kb human *VEGFC-255* RBC region (GRCh37/hg19_dna range: chr4:177,968,343-177,974,845) was sub-cloned into the Kpn1 linearised pGL3-VEGFC promoter-luc-3.4kbHindIII plasmid, with primers (restriction sites were underlined):

*hVEGFC-255* BC/RBC-hP-luc_F (KpnI):

5’-TTTCTCTATCGATA<u>GGTACC</u>AAAGGGTCAAATCGCACAATGC-3’

*hVEGFC-255* RBC-hP-luc_R (KpnI):

5’-TCGCGTAAGAGCTC<u>GGTACC</u>ACAGATTCTCCCAAGTCTGATAATG-3’

Mouse *Vegfc-152* BC-mP-luc, *Vegfc-152* RBC-mP-luc, human *VEGFC-255* BC-hP-luc and *VEGFC-255* RBC-hP-luc constructs were all generated using In-Fusion ® HD cloning (Takara Bio). Fidelity of the inserts were confirmed with sequencing analysis, with primers:

LucNrev: 5’-CCTTATGCAGTTGCTCTCC-3’

RV_Primer3: 5’-CTAGCAAAATAGGCTGTCCC-3’

### Luciferase reporter assay

HeLa cells were seeded in triplicate wells per condition in a 24-well plate at 9x10^4^ cells/well. Culture media was refreshed prior to transfection with 25 ng pRL-TK (Renilla vector purchased from Promega), 250 ng Firefly luciferase plasmids (pGL3-VEGFC promoter-luc-3.4kbHindIII (hP), pGL3-VEGFC promoter-luc-370 (mP), *Vegfc-152* BC-mP-luc, *Vegfc-152* RBC-mP-luc, *VEGFC-255* BC-hP-luc or *VEGFC-255* RBC-hP-luc), 12.5 ng of pCDNA3.1 HEY1 (or the empty vector, pcDNA3.1), or 12.5 ng of pcDNA-SOX7 (or the empty vector, pcDNA3.1 Glomyc) as indicated in figure legends for 24 h, using X-tremeGENE9 DNA transfection reagent according to the manufacturer’s instructions. Cells were washed twice in PBS and harvested using the dual-luciferase assay reporter system, according to the manufacturer’s instructions. To control for transfection efficiency, firefly luciferase activity was determined and normalised to Renilla luciferase activity. Background effects of expression plasmids on pGL3-basic were deducted in all analyses. Readings were then made relative to the human Firefly empty vector-transfected condition, which was set to 1.

### Chromatin immunoprecipitation (ChIP) sequencing and library preparations

Briefly, *Sox7*-V5 embryos were harvested in ice-cold PBS, pooled, minced and digested in 0.05% trypsin/EDTA (Invitrogen) at 37°C for 10 min with constant shaking (5 min for E9.5). Solutions were passed through 21- and 23-gauge needles (5-6 times) to isolate single cells, and cell suspensions were next treated with 1% formaldehyde (for DNA-protein cross-linking) in buffer containing 50 mM Hepes-KOH, 0.1 mM NaCl, 1 mM EDTA, 0.5 mM EGTA for 10 min with constant agitation at room temperature. Subsequently, cell suspensions were quenched with 1 mM glycine pH 7.5 for 5 min at room temperature, washed in ice-cold PBS/10% FBS and finally rinsed in ice-cold PBS. The resultant cell pellet from each litter was snapped frozen on dry ice before storage at -80°C. All steps were carried out on ice unless specified. Cell pellets were next thawed on ice, and pellets across different litters from a similar developmental stage were pooled together before cell lysis. After sonication, 50 μl from each sonication sample was kept aside (this is the input DNA control) before immunoprecipitation step using 10 μg V5 monoclonal antibody (R960CUS, Invitrogen). The subsequent steps were carried out as detailed in the published protocol (Schmidt et al., 2009). Following immunoprecipitation, DNA amplification was performed on both the IP sample and input DNA control using TruSeq ChIPseq kit (Illumina, IP-202-1012) with 0.5 μM of the universal forward and reverse PCR primer containing the index sequence of choice in 50 μL 1 x NEB Next High-Fidelity PCR Master Mix (New England Biolabs, M0541). The number of PCR cycles ranged from 13-18, depending on the ChIP efficiency. The PCR product was purified using AMPure beads (1.8 volume) and eluted in 20 μL of resuspension buffer (Tris-Acetate 10 mM pH 8). The library was quantified using the KAPA library quantification kit for Illumina sequencing platforms (KAPA Biosystems, KK4824) and 50 bp single end reads were sequenced on a HiSeq2500 following the manufacturer’s protocol.

### ChIP-sequencing peak calling, genome annotation and visualisation

After sequencing, FASTA files were uploaded onto Galaxy Australia (Afgan et al., 2015). Quality control was performed on the sequence reads using FastQC (Galaxy version 0.53) (Available from https://www.bioinformatics.babraham.ac.uk/projects/fastqc/). Reads were filtered and trimmed by Trimmomatic (Galaxy version 0.32.2) (Bolger et al., 2014), before alignment to the latest mouse reference genome Genome Reference Consortium (GRC)m38/mm10 (released in Dec 2011) using Bowtie2 (Galaxy version 2.2.6.2) (Langmead and Salzberg, 2012). Repeated sequences were filtered and unique sequences were called using MACS version 2.1.0 (Zhang et al., 2008). Potential binding regions of SOX7 (or peaks) were called using each corresponding input DNA as reference, with a false discovery rate (FDR) of 0.01. The SOX7-V5 ChIP-Seq shows an expected percentage of uniquely mapped reads for a mouse ChIP-seq (Bailey et al., 2013), with >70% unique reads aligned to the reference genome. Only the unique reads were considered for peak calling.

In both SOX7 HUVECs and SOX7-V5 Chip-Seq, binding regions (peaks) were visualised using either UCSC Genome Browser (available at https://genome.ucsc.edu/) or Integrative Genomics Viewer IGV version 2.3.92 (Robinson et al., 2011). Peaks to Genes (top 2k only, ranked by their estimated enrichment folds as depicted in the interval files after peak calling) were annotated by Gene Regions Enrichment of Annotations (GREAT) version 3.0.0 with GRC37/hg19 for human and GRCm38/mm10 for mouse dataset (McLean et al., 2010). Distance of binding regions in relation to the gene transcription start sites (TSS) regions was used as a proxy for the likelihood of transcription regulation.

### *De novo* and local enrichment motif analysis

Motif discovery was performed on the top 2k peaks from E9.5 and E10.5. Sequences in FASTA format 500 bp around the peak centre were created using the “Extract genomic DNA” tool from Galaxy (version 2.2.3) or “GetFastaBed” from Galaxy (version 20.01). The fetched sequences were subsequently piped into Multiple EM for Motif Elicitation (MEME)-ChIP version 5.1.1, using the default settings with the following changes: 1.) ‘JASPAR Vertebrates and UniPROBE Mouse’ was used as the input motif 2.) Under the MEME options, expected motif site distribution was changed to ‘any number of repetitions’, and minimum motif width was changed to 4bp 3.) Under Central Motif Enrichment Analysis (CentriMo), the local mode has been enabled to find un-centred regions (Machanick and Bailey, 2011). Spaced Motif (SpaMo) analysis tool was used to identify enriched spacing between a primary motif and each identified secondary motifs. To identify the total occurrences of the *de novo* motifs discovered by MEME, the Find Individual Motif Occurrences (FIMO) tool was used (Bailey et al., 2009; Bailey et al., 2015). All described tools are available at MEME-suite (http://meme-suite.org/tools/meme-chip).

### Pathway enrichment analysis

Genes associated to the top 2k peaks from SOX7 HUVEC ChIP-Seq (E-MTAB-4480) were used for pathway enrichment search using the GREAT v1.8 Molecular Signatures Database (MSigDB) pathway analysis (Subramanian et al., 2005). Only the top 5 most enriched pathways associated to the gene-set were reported.

### Histone mark and chromatin state SOX7 ChIP-Seq analysis

Bed files with the top 2k of SOX7 HUVEC ChIP-Seq peaks were uploaded onto the EpiExplorer online tool. Histone states and chromatin state segmentation were analysed using the tool function. Regions associated to at least 50% of the chromatin state/ histone markers were then exported into Venny 2.1.0 to generate Venn diagrams. For SOX7-V5 ChIP-Seq, the GRCm38/mm10 version of the top 2k peaks from E9.5 and E10.5 were liftOver to mm9 using the UCSC Genome Browser. These converted tracks were then uploaded on the EpiExplorer online tool to be overlapped with histone markers from the ENCODE database. Chromatin State Segmentation analysis is not available for mouse data. Corresponding regions associated to at least 50% overlapping with each histone marks were next exported into Venny 2.1.0 to generate Venn diagrams.

### Antibodies

For immunostaining, primary antibodies were: rat anti-EMCN (1:300, sc-53941, Santa Cruz), goat anti-NRP2 (1:300, AF567, R&D System), goat anti-SOX7 (1:300, AF2766, R&D System), goat anti-SOX17 (1:300, AF1924, R&D System), rabbit anti-PROX1 (1:300, 11-002, AngioBio), rabbit anti-ERG (1:300, AB92513, Abcam), rat anti phospho-histone H3 (1:500; H9908, Sigma Aldrich), rabbit anti-V5 (1:200, AB3792, Merck), chicken anti-beta galactosidase (1:200, AB9361, Abcam), DAPI (1:1000, D9542, Sigma Aldrich).

Secondary antibodies were: goat anti-chicken IgG Alexa 488 (A11039), donkey anti-goat IgG Alexa 647 (A21447), donkey anti-rat IgG Alexa 488 (A21208), donkey anti-rabbit Alexa 594 (A21207), donkey anti-rabbit IgG Alexa 647 (A31573), donkey and anti-goat IgG Alexa 594 (A11058), goat anti-rat IgG Alexa 594 (A11007), goat anti-rat IgG Alexa 647 (A21247). Secondary antibodies were sourced from Invitrogen and used at 1:300 unless specified otherwise.

For co-immunoprecipitation and western blotting, primary antibodies were: Living Colors Ds-red rabbit antibody (632496, Clontech), rabbit anti-tubulin (T3526, Sigma-Aldrich), and mouse anti-GFP (3E6) (A-11120, Thermo-Scientific). Secondary antibodies were: goat anti-rabbit IgG HRP (65-6120), goat anti-mouse IgG HRP (A10551). Secondary antibodies were sourced from Invitrogen, all antibodies were used at 1:1000.

For cell-sorting experiment, antibodies were: Alexa 488 conjugated rat anti-CD31 (1:200, 563607, BD Pharmigen), PE/Cy5 rat anti CD45 (1:1000, 103109, BioLegend), live/dead cells were detected by DAPI (1:1000, D9542, Sigma Aldrich).

### cDNA synthesis and real-time PCR

First strand cDNA was synthesised from 1500 ng purified RNA using High-Capacity cDNA Reverse Transcription Kit (Life technologies). qPCR was performed and analysed as described in (Chiang et al., 2017). To quantify the transcript level of target genes, primers used were:

mSox7_F: 5’-GCGGAGCTCAGCAAGATG-3’

mSox7_R: 5’-GGGTCTCTTCTGGGACAGTG-3’

mSox17 _F: 5’-CACAACGCAGAGCTAAGCAA-3’

mSox17_R: 5’-CGCTTCTCTGCCAAGGTC-3’

mSox18 _F: 5’-ACTGGCGCAACAAAATCC-3’

mSox18_R: 5’-CTTCTCCGCCGTGTTCAG-3’

mVegfr3_F: 5’-GGTTCCTGATGGGCAAAGG-3’

mVegfr3_R: 5’-TCAGTGGGCTCAGCCATAGG-3’

mHey1_F: 5’-CATGAAGAGAGCTCACCCAGA-3’

mHey1_R: 5’-CGCCGAACTCAAGTTTCC-3’

mVegrf2 F: 5’-CAGTGGTACTGGCAGCTAGAAG-3’

mVegfr2_R: 5’-ACAAGCATACGGGCTTGTTT-3’

mDll4_F: 5’-AGGTGCCACTTCGGTTACAC-3’

mDll4_R: 5’-GGGAGAGCAAATGGCTGATA-3’

mNotch1_F: 5’-CTGGACCCCATGGACATC-3’

mNotch1_R: 5’-AGGATGACTGCACACATTGC-3’

mPecam_F: 5’-CGGTGTTCAGCGAGATCC-3’

mPecam_R: 5’-ACTCGACAGGATGGAAATCAC-3’

mCdh5_F: 5’-GTTCAAGTTTGCCCTGAAGAA-3’

mCdh5_R: 5’-GTGATGTTGGCGGTGTTGT-3’

mVegfc primers sequenced were from (Hominick et al., 2018).

hSOX7_F: 5’-AGCTGTCGGATGGACAATCG-3’

hSOX7_R: 5’-CCACGACTTTCCCAGCATCT-3’

hVEGFR3_F: 5’-ATAGACAAGAAAGCGGCTTCA-3’

hVEGFR3_R: 5’-CCTCCCTTGGGAGTCAGG-3’

hVEGFR2_F: 5’-GAACATTTGGGAAATCTCTTGC-3’

hVEGFR2_R: 5’-CGGAAGAACAATGTAGTCTTTGC-3’

hGAPDH_F: 5’-CCCCGGTTTCTATAAATTGAGC-3’

hGAPDH_R: 5’-CACCTTCCCCATGGTGTCT-3’

hHEY1_F: 5’-CATACGGCAGGAGGGAAAG-3’

hHEY1_R: 5’-GCATCTAGTCCTTCAATGATGCT-3’

hVEGFC_F: 5’-CACTACCACAGTGTCAGGCA-3’

hVEGFC_R: 5’-GTCATCTCCAGCATCCGAGG-3’

hVEGFA_F: 5’-AGGGAAAGGGGCAAAAACGA-3’

hVEGFA_R: 5’-CCTCGGCTTGTCACATCTGC-3’

hPROX1_F: 5’-TCACCTTATTCGGGAAGTGC-3’

hPROX1_R: 5’-GAGCTGGGATAACGGGTATAAA-3’

hGAPDH_F: 5’-CCCCGGTTTCTATAAATTGAGC-3’

hGAPDH_R: 5’-CACCTTCCCCATGGTGTCT-3’

hHPRT_F: 5’-AATGACCAGTCAACAGGGGACA-3’

hHPRT_R: 5’-TACTGCCTGACCAAGGAAAGCA-3

### Quantification and data analysis

To quantify the proportion of SOX7 positive cells in Fig. 1A, SOX7+ cells were calculated manually using ImageJ, and subsequently divided by ERG+ endothelial cells in the entire section. Fig. 1D-J were quantified within 2100 μm from both sides of the midline. Migration distance (Fig. 1D) were an average of 10 random measurements taken from the midline to the nearest leading lymphatic sprouts. For vessel width (Fig. 1F), 3 locations within 200 μm from the sprouting end of each lymphatic leading vessel was measured. Vessel width was taken from the average of 7 representative lymphatic leading vessels in each embryo. PROX1+ nuclei in Fig. 1H were an average of PROX1+ number quantified from the 7 representative lymphatic leading vessels in Fig. 1F (200 μm from the sprouting end). For Fig. 1I, disconnected vessels (>100 μm) or lymphatic endothelial cell (LEC) clusters (<100 μm) are defined as PROX1+ NRP2+ cell population that is isolated from the lymphatic network. Only the PROX1+ nuclei in single LEC cluster were quantified and shown in Fig. 1J.

To quantify the % of PROX1+ cells within indicated region in the cardinal veins (CVs) (Fig. 5B), CVs were divided into a 12-section pie chart. Region nearest to the dorsal aorta was designated as 0/360°. Within each region, number of PROX1+ LEC progenitors and EMCN+DAPI+ endothelial cells were quantified. Each region from Fig. 5A-C were the average of 20 and 17 vibratome sections from sibling controls (n=5) and *Sox7^iECKO^* mutants (n=4), respectively. Only sections from middle to lower thoracic regions were included for analysis. Data were then plotted in a rose diagram using Visual Paradigm online Diagrams.

To quantify the number of PROX1+ LECs or LEC clusters associated to EMCN+ BV (Fig. 5D-E), confocal images were taken randomly from each skin quadrants: left/ right cervical, thoraco-cervical and thoraco-lumbar regions. To effectively quantify the PROX1+ LECs associated to LV, EMCN+ surface was created by the “surface” function in Imaris, which was then masked for PROX1+ cells. Then, PROX1+ cells were counted manually by carefully going through each image slices, and scrutinised also in 3D projection. PROX1+ cells which belongs to the greater lymphatic networks and isolated from BV were excluded. PROX1+ cell-type were categorised according to Sup. Fig. 8A, B. Where a PROX1+ cell or cluster appears to be associated to BV, a confocal image at higher resolution (X40-X63) would be taken for further confirmation. Total PROX1+ cells were finally normalised to the total BV surface area analysed.

Vessel front density (Sup. Fig. 4B) was quantified from EMCN+ blood vessels 500 μm from the midline (700 x 3072 μm) using the “surface” function in Imaris. The number of H3+ERG+ proliferative endothelial cells and total EGR+ endothelial cells from the same region were quantified with the “spot” function in Imaris.

Sup. Fig. 4D-F were quantified from a similar area of skins (3500 x 2000 μm), using the “surface” and “spots” function in Imaris. Total blood vessel (BV) surface density was surface rendering of the EMCN+ vessel (Sup. Fig. 4D). Green BV was green area that co-localised with EMCN+ surface. Percentage of Cre recombination in BV was therefore green BV density / total BV density. Total lymphatic endothelial cells (LECs) were quantified by spots rendering of PROX1+ nuclei (Sup. Fig. 4E). Total green LECs were determined by spots rendering of green cells, which were also PROX1+. The percentage of Cre recombination in lymphatic vessel (LV) was therefore Green LECs/Total LECs. BV blood density was total EMCN+ BV surface (Sup. Fig. 4F).

## ACKNOWLEDGEMENT

The authors would like to thank Dr Ming-Jen Hsu for the kind gift of the mouse luciferase reporter construct, pGL4-VEGFC promoter-luc-370, Dr Emma Gordon for the *mTmG* mice, Dr Luciano Martelotto and Dr Yen Tran for nuclei isolation protocol in single-nuclei RNA Seq. We thank Dr Carol Wicking for scientific comments and advice on editing of the manuscript.

## Author contributions

IC and MF were involved in the design of the experiments. IC, WL, NK, MM, RS, and TD performed experiments and collated data. IC, WL, KY, NK, MM, and MF were involved in data analyses. IC and MF wrote the manuscript constructed the figures. All authors commented on the manuscript.

## Supplementary Figures

**Supplementary Figure 1.**
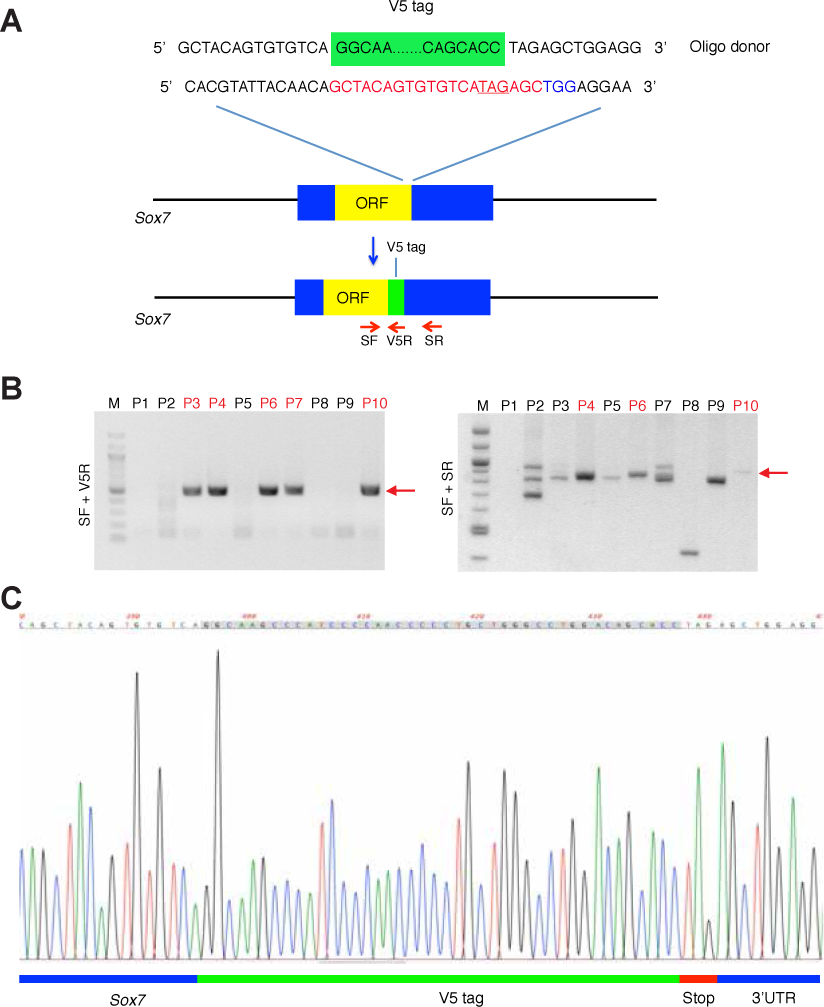
Schematic showing the CRISPR design to insert the V5 tag into the Sox7 locus. (A) Cas9/sgRNA/Oligo-targeting site near the SOX7 stop codon (underlined in red). The SgRNA sequence is highlighted in red (this contains the stop codon), while the protospacer-adjacent motif (PAM) sequence is shown in blue. The oligo donor containing the V5 tag (green box), is flanked by 60 bp homology arms. The red arrows represent PCR sequencing primers for genotyping (SF, V5R and SR). **(B)** PCR genotyping of CRISPR *Sox7*-V5 injected live-born pups. Left: PCR genotyping with primer pair SF and V5R showed the desired band at the correct size in postnatal (P) 3, P4, P6, P7 and P10. Right: PCR genotyping with primer pair SF and SR produced slightly larger products in P4, P6 and P10 compared to P3 and P7, indicating successful 42 bp V5 tag integration. At least 3 bands were discovered in P3 and P7, but not in P4, P6 and P10 when separated by 3.5% gel electrophoresis (not shown). Red arrows indicate the expected PCR product sizes. **(C)** Sequencing of PCR product using SF primer (for both SF/V5R and SF/SR) confirmed the correct fusion of V5 tagged to the last codon in both *Sox7* alleles of the *Sox7*-V5 founders.

**Supplementary Figure 2.**
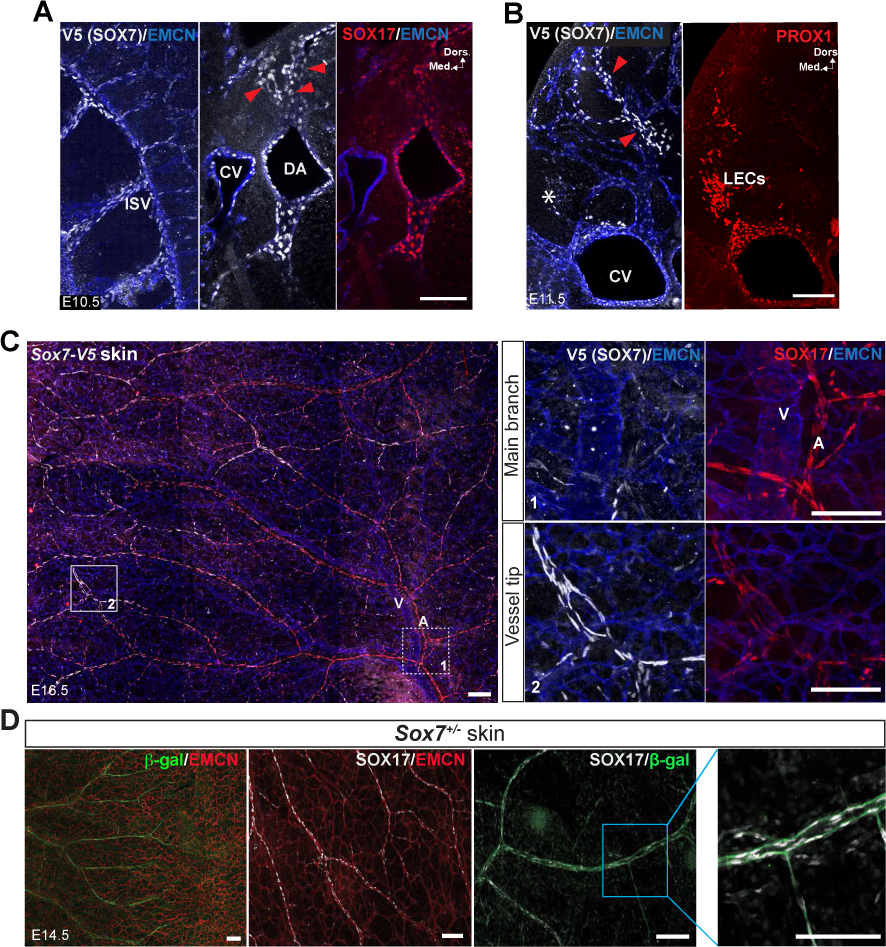
SOX7-V5 expression recapitulates endogenous SOX7 profile. (A) At E10.5, SOX7 (white) is expressed in intersomitic vessels (ISVs), the dorsal aorta (DA) (delineated by arterial marker, SOX17), cardinal vein (CV), and the surrounding migrating endothelial cells (red arrowheads) **(B)** SOX7 was not detected in PROX1+ migrating lymphatic endothelial cells (LECs) (asterisks). **(C)** At E16.5, SOX7 is downregulated in the SOX17+ main arterial branch (boxed area with dotted line), but continues to be expressed in the less mature arteries near the midline (box area with solid line). Arteries are delineated by arterial marker SOX17; major veins and vascular plexus are delineated by endomucin (EMCN). **(D)** SOX7/*β*-galactosidase (*β*-gal) (green) also stains the SOX17 (white) positive major arteries in the skin plexus of E14.5 *Sox7^+/-^lac-Z* reporter mice. Dors., dorsal; Med., medial. Scale bars = 100 μm.

**Supplementary Figure 3.**
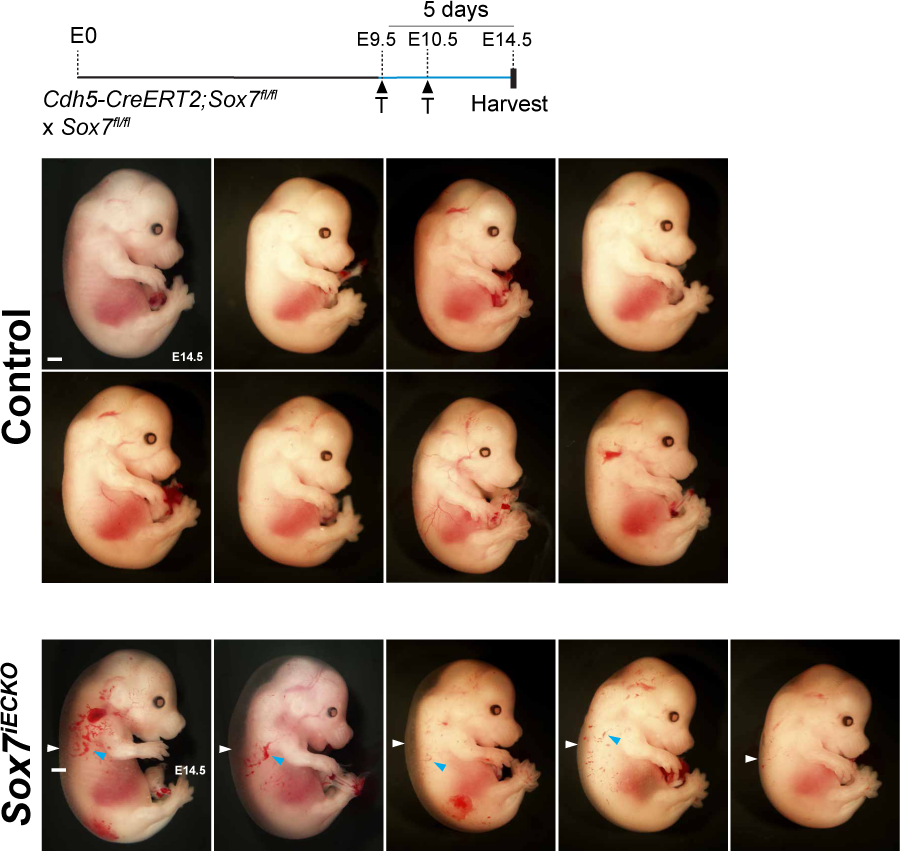
Brightfield images of *Sox7^iECKO^* mutants and sibling controls. Embryos are collected at E14.5, after pulsed with tamoxifen at E9.5 and E10.5. All mutants developed subcutaneous edema (white arrowheads), with the 4 of them showing blood-filled lymphatics (blue arrowheads). Scale bars = 1 mm.

**Supplementary Figure 4.**
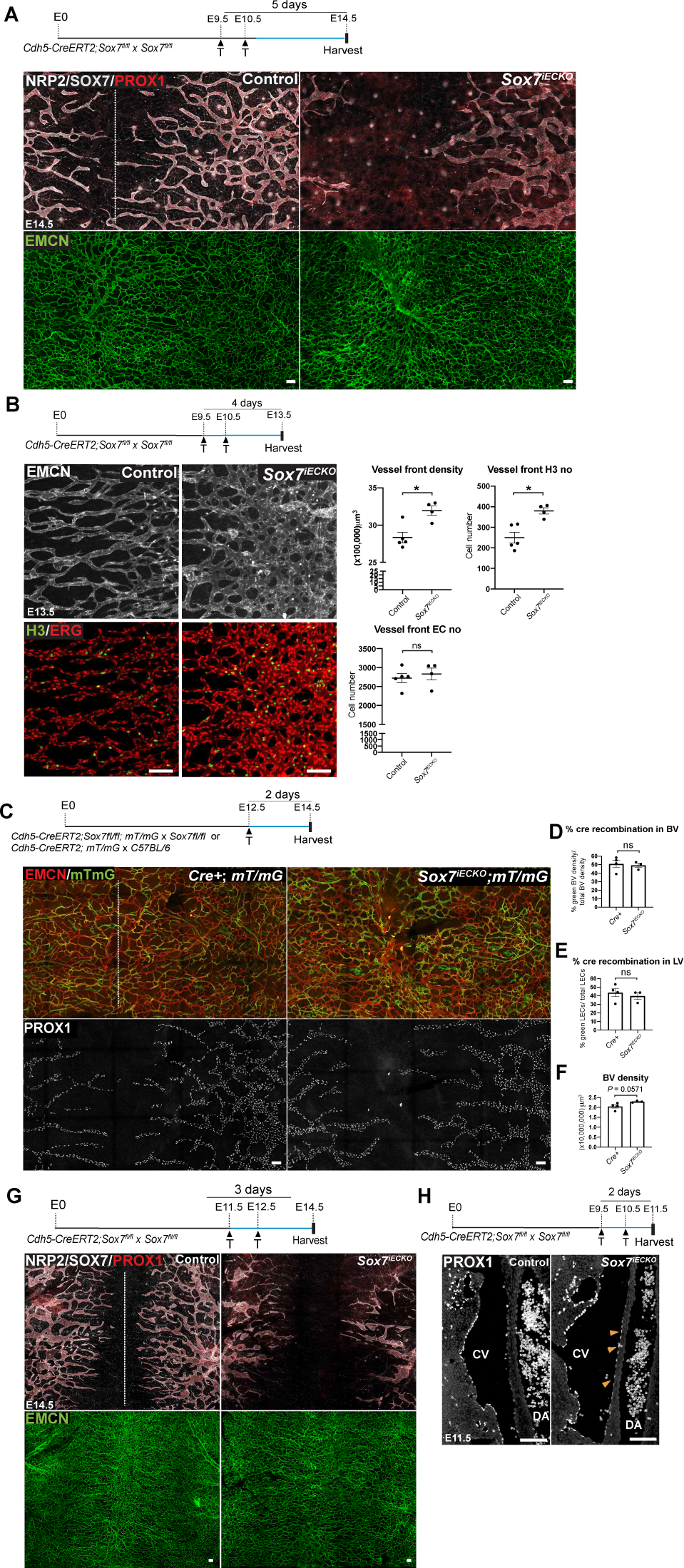
Vascular defects associated with loss of SOX7 function. (A) Whole-mount immunostaining of *Sox7^iECKO^* mutant and sibling control embryonic skin at E14.5, following injection with tamoxifen at E9.5 and E10.5. Dermal lymphatic structures are marked by NRP2 (membranous white), lymphatic endothelial cells by PROX1 (red), blood vessels by EMCN (green) and arterioles by SOX7 (white nuclei). Dashed line represents the midline of the embryo. **(B)** Whole-mount immunostaining of *Sox7^iECKO^* mutant and sibling control for EMCN (grey), showing dermal blood vessels at E13.5, after injection with tamoxifen at E9.5 and E10.5. Endothelial cells are marked by ERG (red), proliferative cells are stained by phospho-histone 3, H3 (green). (Right panel) Quantitation of EMCN+ (blood) vessel front density (μm^3^), number of H3+ proliferative and ERG+ endothelial cells in this region. Scored sibling control, n=5; *Sox7^iECKO^* mutants, n=4; Mean ± SEM; Mann-Whitney *U*-test. *P*<0.05 (*); ns = not significant. **(C)** Whole-mount immunostaining of *Sox7^iECKO^*; *mT/mG* mutant and *Cre+; mT/mG* control embryonic skin at E14.5, after injection with tamoxifen at E12.5. Blood vessels are stained with EMCN (red), cells after *Cre* excision are shown in green, and lymphatic endothelial cells are marked by PROX1 (white). **(D-F)** Graphs showing quantitation of the Cre activity in (D) blood vessels, (E) lymphatic vessels and (F) EMCN+ (blood) vessel density (μm^3^), from n=4 *Cre+; mT/mG* controls and n=3 *Sox7^iECKO^*; *mT/mG* mutants. Mean ± SEM; Mann-Whitney *U*-test; ns = not significant. **(G)** Whole-mount immunostaining of *Sox7^iECKO^* mutant and sibling control embryonic skin at E14.5, after injection with tamoxifen at E11.5 and E12.5. Dermal lymphatic structures are marked by the NRP2 (membranous white), lymphatic endothelial cells by PROX1 (red), and blood vessels are stained by EMCN (green). Dashed line represents the midline of the embryo. **(H)** Immunostaining of *Sox7^iECKO^* mutant and sibling control coronal sections, at E11.5, after injection with tamoxifen at E9.5, E10.5. Lymphatic progenitor cells in the cardinal veins (CVs) are PROX1+ (white). Yellow arrowheads highlight the presence of LEC progenitors in the region near the dorsal aorta (DA) in the *Sox7^iECKO^* mutant, which is typically absent in the sibling control. Scale bars = 100 μm.

**Supplementary Figure 5.**
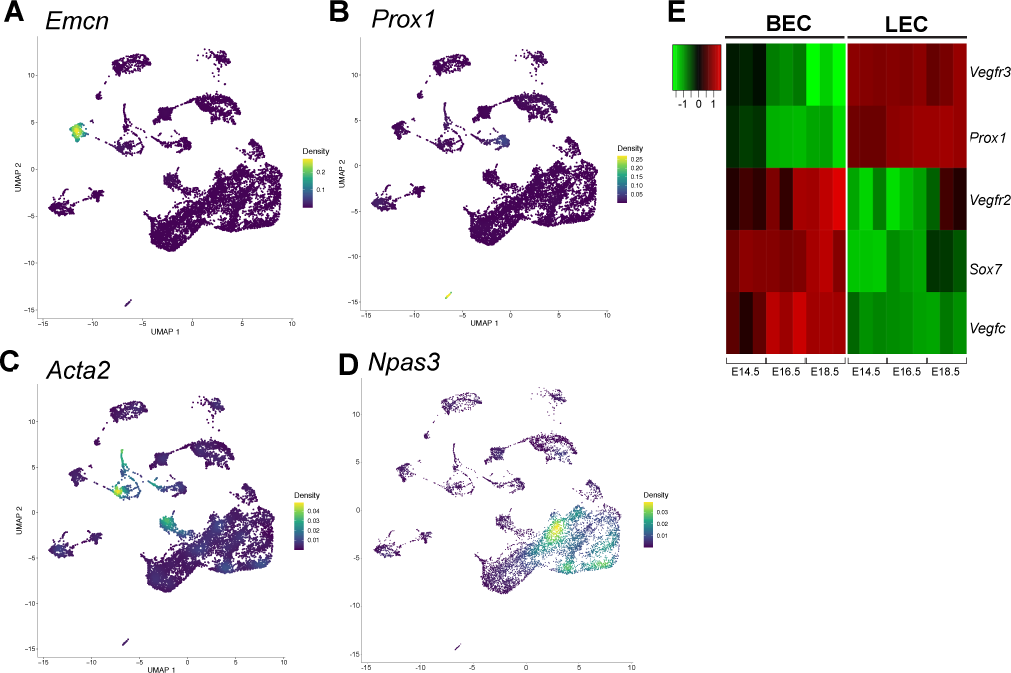
(A-D) Nebulosa plots from single-nuclei RNA-Seq on E14.5 embryonic skins, showing markers for BECs (*Emcn*), LECs (*Prox1*), SMCs (*Acta2*) and neuronal cells (*Npas3*). **(E)** Microarray analysis of dermal BEC and LEC populations sorted from different embryonic stages reveals that *Sox7* and *Vegfc* mRNA are preferentially expressed by BECs. BECs, blood endothelial cells; LECs, lymphatic endothelial cells; and SMC, smooth muscle cells.

**Supplementary Figure 6.**
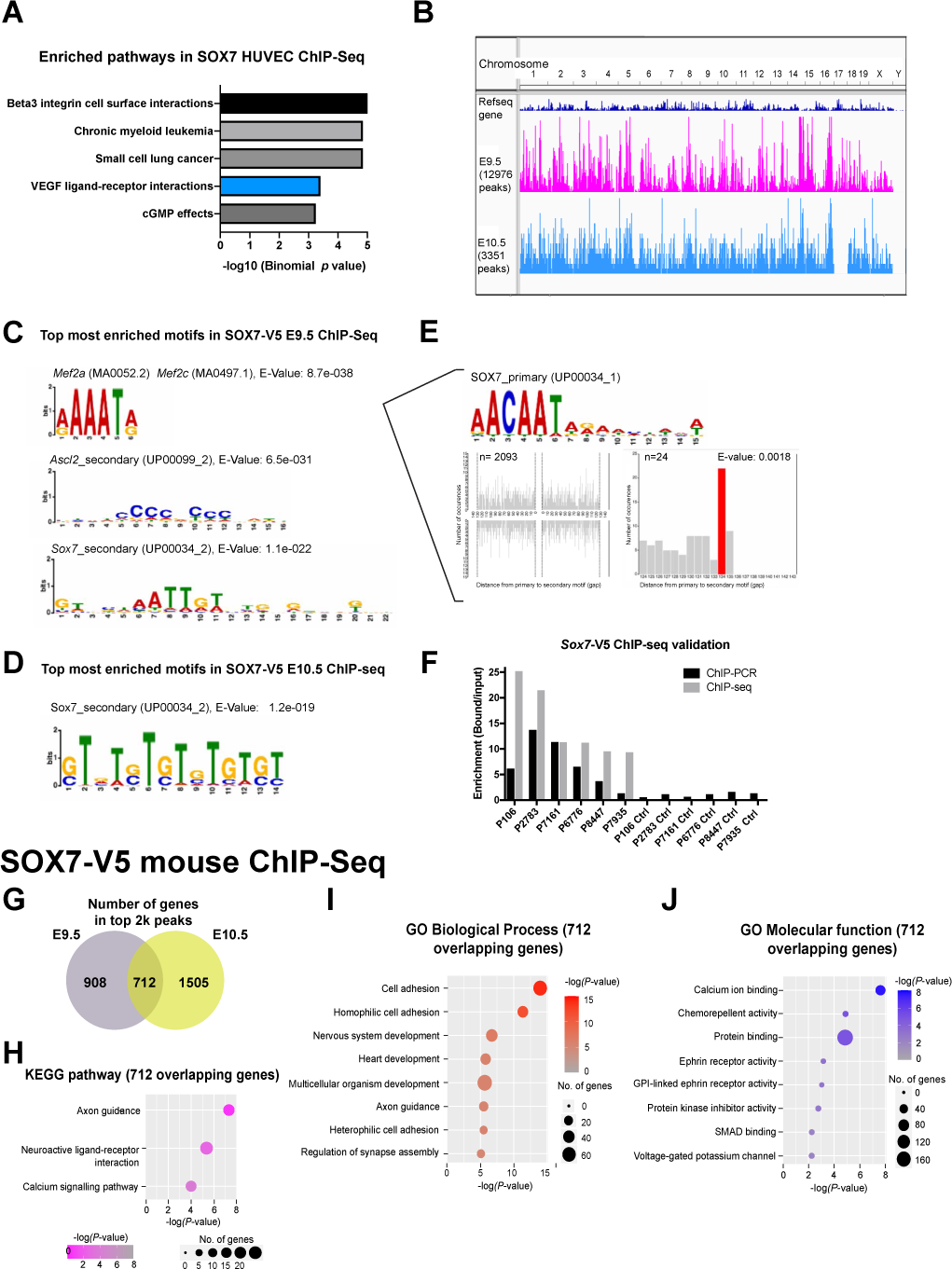
Downstream analyses of SOX7 ChIP-Seq. (A) MSigDB pathway analysis on the top 2k peaks from human SOX7 HUVEC ChIP-Seq identified enrichment for genes within the VEGF/VEGFR gene set. These include *VEGFR1*, *VEGFR2*, *NRP1*, *NRP2*, *PDGFC*, *VEGFA* and *VEGFC*. **(B-F) Binding profiles and motif analysis of mouse SOX7-V5 ChIP-Seq (B)** Image from Integrative Genome Viewer (IGV) illustrating ChIP-Seq tracks at E9.5 (pink) and E10.5 (light blue) mapped to the mouse reference genome GRCm38/mm10 (top track, dark blue). The height of a graph peak indicates the number of “positive bindings” on a given chromosomal location. Total number of binding events is indicated on the left under each track name. Top DNA-binding motifs in SOX7*-*V5 ChIP-Seq by MEME-ChIP in **(C)** E9.5 and **(D)** E10.5 embryos. The different height of the letters in the position weight matrix reveals information content of each position (in bits), correlated by the degree of certainty of the nucleotide at a given position. **(E)** Spaced motif analysis (SPAMO) found a SOX7 primary motif amongst the topmost enriched motif next to a MEF2A/MEF2C binding motif found in E9.5 SOX7*-*V5 ChIP-Seq, with 2093 total occurrences. SOX7 is best gapped at 134 bp from the MEF2A/MEF2C binding motif (24/2093). **(F)** ChIP-PCR on both bound fraction (left) and corresponding assumed unbound controls (right), as predicted by the peak location from SOX7-V5 ChIP-Seq. **(G)** A total of 712 genes were found overlapping between E9.5 and E10.5 SOX7-V5 mouse ChIP-Seq. **(H-J)** KEGG pathway analysis **(H)** and gene ontological analysis by DAVID bioinformatics database, showing the top biological processes **(I)** and molecular functions **(J)** associated to the 712 overlapping genes from the SOX7-V5 mouse ChIP-Seq.

**Supplementary Figure 7.**
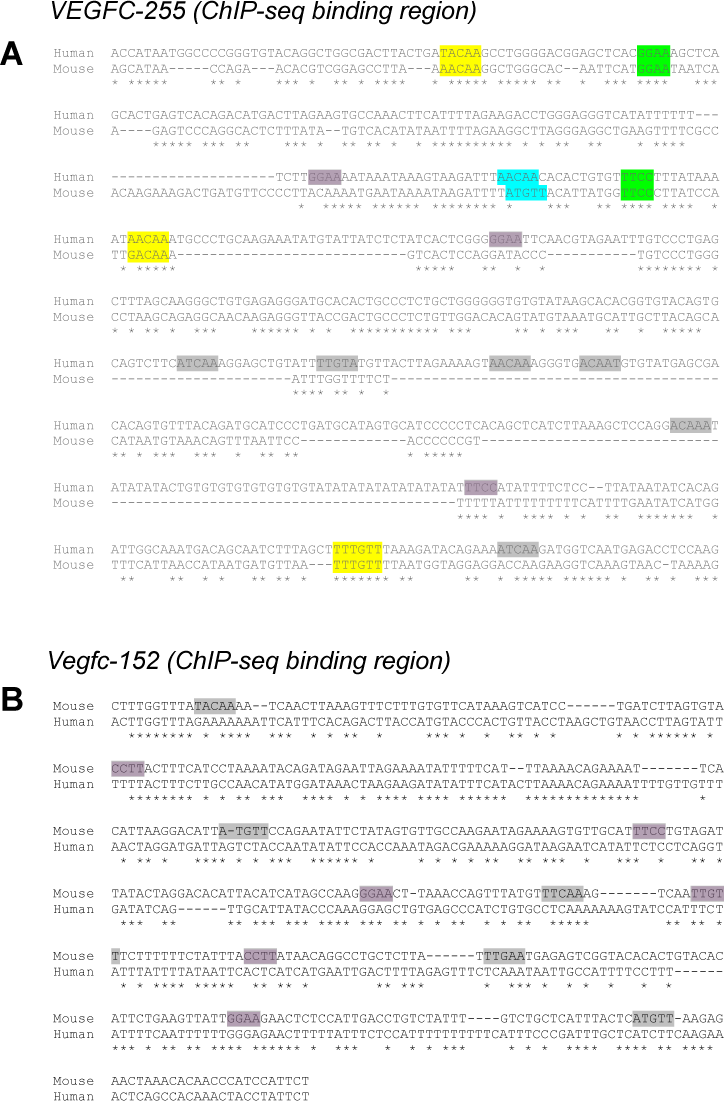
Multiple species alignment of the orthologous regions of the (A) *VEGFC-255* and (B) *Vegfc-152* binding region from human and mouse. Yellow and green boxes indicate conserved regions across human and mouse for SOX and ETS, respectively. Cyan box indicates conserved SOX motif (different nucleotide bases). Grey (SOX) and purple (ETS) boxes indicate un-conserved regions. Alignment was performed using ClustalW.

**Supplementary Figure 8.**
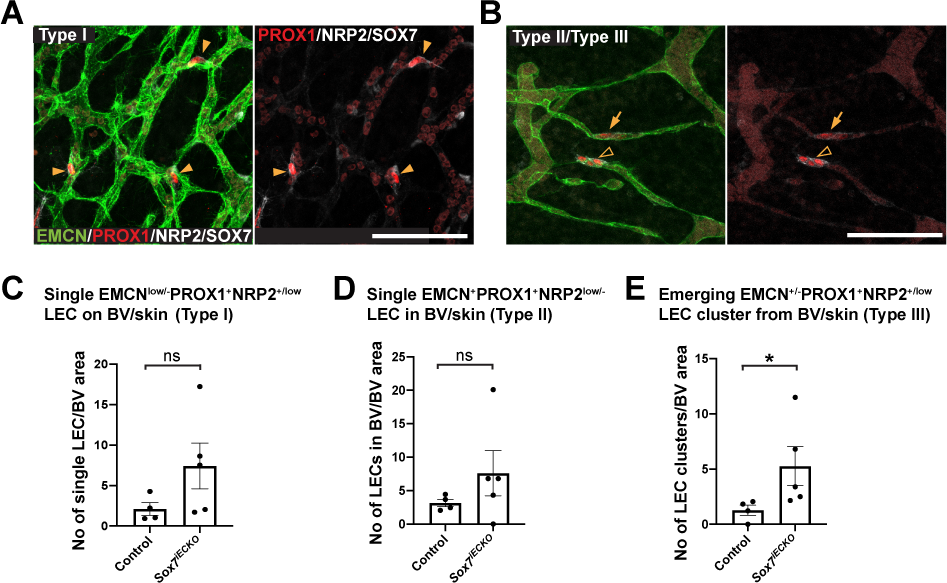
Loss of SOX7 increases the number of LEC progenitors specified in blood plexus. (A-B) Whole-mount immunostaining of *Sox7^iECKO^* mutant or control embryonic skin, showing the different types of LECs emerging from the blood capillary plexus. Skins were harvested from embryos at E12.5, after Cre-induction at E9.5 and E10.5. Dermal lymphatic structures are marked by NRP2 (membranous white), lymphatic endothelial cells by PROX1 (red), and blood vessels are stained by EMCN (green) and SOX7 (white nuclei). Arrowheads indicate EMCN^low/-^PROX1^+^NRP2^+/low^ individual LECs on blood vessels (type I), arrows show the single EMCN^+^PROX1^+^NRP2^low/-^ LEC within the blood vessel wall (type II) and empty arrowheads mark the EMCN^+/-^PROX1^+^NRP2^+/low^ LEC clusters (2-4 cells) exiting from the blood vessel network (type III). **(C-E)** Graph shows the quantitation of different types of PROX1+ LEC progenitors, comparing between control and *Sox7^iECKO^* mutant skins. PROX1+ LEC progenitors were quantified from n=4 controls and n=5 *Sox7^iECKO^* mutants. Mean ± SEM; Mann-Whitney *U*-test. *P*<0.05 (*); ns = not significant. Scale bars = 100 μm.

**Supplementary Figure 9.**
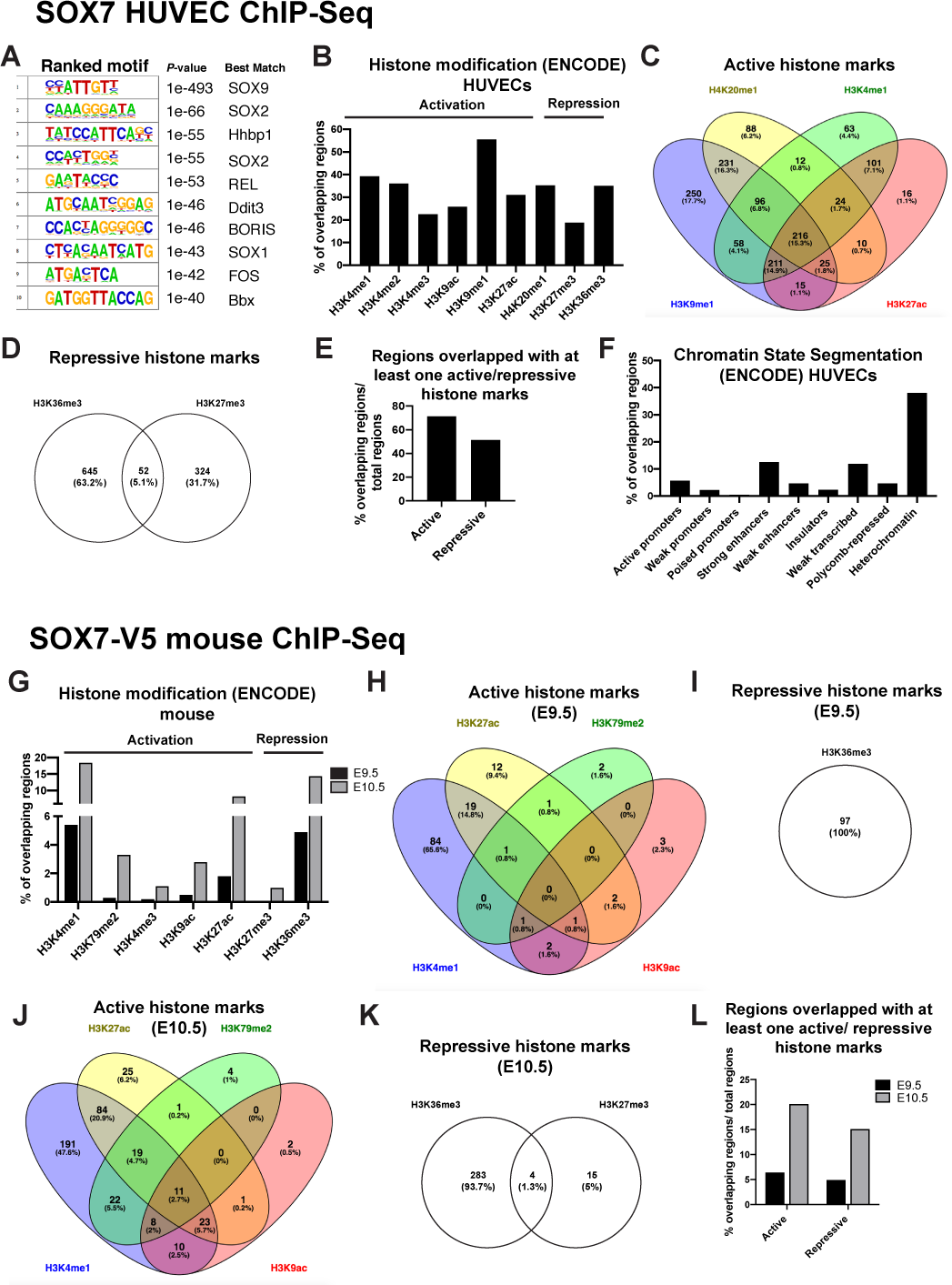
*In silico* characterisation of SOX7 HUVECs and SOX7-V5 ChIP-Seq. (A-F) SOX7 HUVEC ChIP-Seq. (G-L) SOX7-V5 ChIP-Seq. (A) Table from HOMER motif database showing the top 10 most enriched transcription motifs associated to SOX7 binding sites **(B, G)** Graphs showing intersections between top 2k SOX7 ChIP-Seq peak locations and HUVEC or mouse histone markers, respectively from the ENCODE consortium. Only peaks with at least 50% overlapping region with the histone markers are considered. Intersection was performed using the EpiExplorer online resource tool. **(C, H, J)** Venn diagrams showing intersections of different active histone markers and **(D, I, K)** repressive histone markers, both displayed at least 50% overlapped, with the top 2k of SOX7 ChIP-Seq binding regions. Diagram was generated using online resource stool, Venny 2.1.0. **(E, L)** Graphs showing the overall proportion of top 2k SOX7 ChIP-Seq peaks, with at least an active or a repressive histone mark. **(F)** Graphs showing intersections between top 2k SOX7 HUVEC ChIP-Seq peak locations and chromatin states in HUVECs from the ENCODE consortium. Only peaks with at least 50% overlapping region with each corresponding chromatin state are considered. Intersection was performed using the EpiExplorer online resource tool.

**Sup. Table 1.**
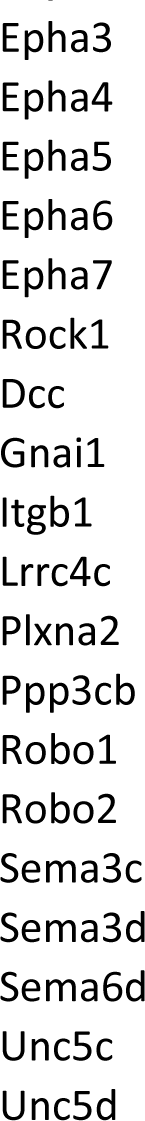
SOX7-V5 ChIP-seq genes related to “Axon Guidance” in KEGG pathway

